# SNAC-DB: An ML-Ready Database for Antibody and NANO-BODY® VHH–Antigen Complexes with Expanded Structural Diversity and Real-World Benchmarking

**DOI:** 10.64898/2026.04.22.720253

**Authors:** Abhinav Gupta, Bryan Munoz Rivero, Ruijiang Li, Jorge Roel-Touris, Yves Fomekong Nanfack, Maria Wendt, Yu Qiu, Norbert Furtmann

**Author notes:** Corresponding Authors: Abhinav Gupta, Large Molecule Research, Sanofi, 350 Water St, Cambridge, MA 02141, United States, Norbert Furtmann, Large Molecule Research, Sanofi, Industriepark Höchst, 65926 Frankfurt am Main, Germany. These authors contributed equally to this work (Abhinav Gupta, Bryan Munoz Rivero). Work performed during co-op placement at Sanofi.

## Abstract

Predicting antibody and NANOBODY^®^ VHH–antigen complexes remains a critical challenge for state-of-the-art structure prediction models, limiting their impact in therapeutic discovery pipelines. We introduce SNAC-DB, an ML-ready database and curation pipeline enriched with structural biology expertise, designed to accelerate model accuracy and generalization by providing 31–37% expanded structural diversity over existing resources like SAbDab through comprehensive re-curation that extracts maximum value from available experimental structures. SNAC-DB expands coverage by capturing often-overlooked complexes and accurately identifying complete multi-chain epitopes through improved biological-assembly-based logic. Built for ML practitioners, SNAC-DB provides standardized formats with multi-threshold structure-based clustering to enable principled sample weighting during training. Using a rigorous benchmark of public PDB entries deposited post-May 2024 plus confidential therapeutic structures, we evaluate seven leading models (Protenix-v1, OpenFold-3p2, RosettaFold-3, Boltz-2, Boltz-1x, Chai-1, and AlphaFold2.3-multimer) with evaluation methodology tailored to antibody/NAN-OBODY^®^ VHH–antigen complexes to ensure correct handling of multi-chain epitopes, revealing systematic performance gaps: success rates rarely exceed 25%, confidence-based ranking fails to identify best predictions even when accurate structures exist in ensembles, and all models consistently struggle with therapeutically relevant NANOBODY^®^ VHHs. Systematic evaluation of sampling strategies demonstrates that while generating 1000 samples per target substantially increases the likelihood of producing accurate structures (oracle selection improves from 11.9% to 50.5%), confidence-based ranking remains nearly flat (between 10.9% and 14.9%), revealing that improved ranking mechanisms represent a more tractable path to performance gains. Finally, fine-tuning GeoDock on SNAC-DB yields higher success rates than training on SAbDab (11.0% vs. 7.1% for antibodies; 7.0% vs. 4.0% for NANOBODY^®^ VHHs), suggesting that SNAC-DB’s expanded structural diversity translates to improved model generalization.

**Significance Statement:** Computational antibody/NANOBODY^®^ VHH design shows promise but remains unreliable for therapeutic development. SNAC-DB provides 31–37% expanded structural diversity through comprehensive data curation, immediately accelerating model development. Benchmarking seven leading AI models reveals accuracy rarely exceeds 25% on therapeutic targets, with confidence-based ranking failing to identify correct structures even when they exist in model outputs. Training on SNAC-DB increases prediction accuracy, validating that high-quality, diverse training data is critical for advancing computational methods toward clinical impact.

## 1 Introduction

Over the past few decades, the Protein Data Bank (PDB) (Berman et al., 2000) has underpinned Nobel Prize-winning breakthroughs in structural biology, including AlphaFold (Jumper et al., 2021b) and computational protein design (Huang et al., 2016), while fueling the rise of AI/ML and classical bioinformatics pipelines. These advances have dramatically enhanced our ability to predict and engineer protein–protein interactions, yet antibody (Ab) and NANOBODY^®^ VHH (Nb)–antigen (Ag) complexes—restricted here to protein antigens—remain a persistent blind spot. Despite optimistic benchmarking reports, cutting-edge complex prediction methods frequently misidentify epitopes and fail to recover key interface contacts (Hitawala and Gray, 2024; Yin and Pierce, 2024). In practice, Ab/Nb–Ag modeling still trails general protein–protein prediction, which can leverage coevolutionary signals from paired multiple sequence alignments (MSAs) and typically involves relatively rigid interfaces. In contrast, complementarity-determining regions (CDR loops) are hypervariable, highly flexible, and often undergo conformational rearrangements upon binding, making accurate prediction substantially more challenging.

For over a decade, the Structural Antibody Database (SAbDab) (Dunbar et al., 2014) has served as the *de facto* source for curated Ab/Nb complexes, with weekly updates powering everything from homology-based modeling to large-scale deep learning. However, raw PDB files remain notoriously complex: chain identifiers vary, biological assemblies and asymmetric units may be mislabeled or incomplete, missing residues are recorded only in headers, and true binding partners can be obscured by crystal-packing artifacts or multi-chain epitopes. While SAbDab applies rule-based parsing to extract complexes, such automated approaches inevitably miss edge cases that require expert structural biology insight. Moreover, SAbDab’s strict focus on Ab/Nb interactions with non-immunoglobulin targets excludes several classes of therapeutically relevant complexes—most notably Ab–Nb pairs and TCR–antigen assemblies—that are increasingly important for rational therapeutic design. Although high-quality TCR–antigen structures are available from specialized resources like TCR3d (Gowthaman and Pierce, 2019), most immunoglobulin-centric ML pipelines overlook these functionally informative assemblies.

Current ML workflows also face common preprocessing pitfalls. The prevalent practice of redundancy filtering—typically based on arbitrary sequence-identity or resolution thresholds (e.g., excluding structures with resolution *>* 4 Å)—can inadvertently remove structures capturing novel targets or epitopes, omit variants differing by a few functionally critical mutations, and discard multiple binding modes on the same antigen. Furthermore, existing repositories provide data in formats optimized for structural biologists rather than ML practitioners: cryptic chain labels for synthetic constructs, missing-residue information confined to PDB headers or FASTA files, and the legacy PDB versus mmCIF format choice all force users to write custom extraction routines. Converting coordinates into ML-friendly representations (e.g., NumPy arrays in *atom37* format introduced in the AlphaFold2 codebase (Jumper et al., 2021a)) further compounds the risk of silent preprocessing errors that can compromise model training.

In this study, we introduce SNAC-DB (*Structural NANOBODY*^*®*^ *VHH and Antibody Complex Database*), a comprehensive ML-ready resource designed to address these limitations. SNAC-DB provides automated curation of Ab–Ag, Nb–Ag, and TCR–antigen complexes from PDB biological assemblies, accurate multi-chain epitope detection, inclusion of often-overlooked complex types, and delivery in standardized formats with transparent structure-based clustering for principled sample weighting. Through comprehensive re-curation, SNAC-DB achieves up to 37% expanded structural diversity over SAbDab, enabling researchers to extract maximum value from available experimental data without waiting for future structure deposits. Figure 1 provides an overview of the SNAC-DB curation pipeline.

**Figure 1:**
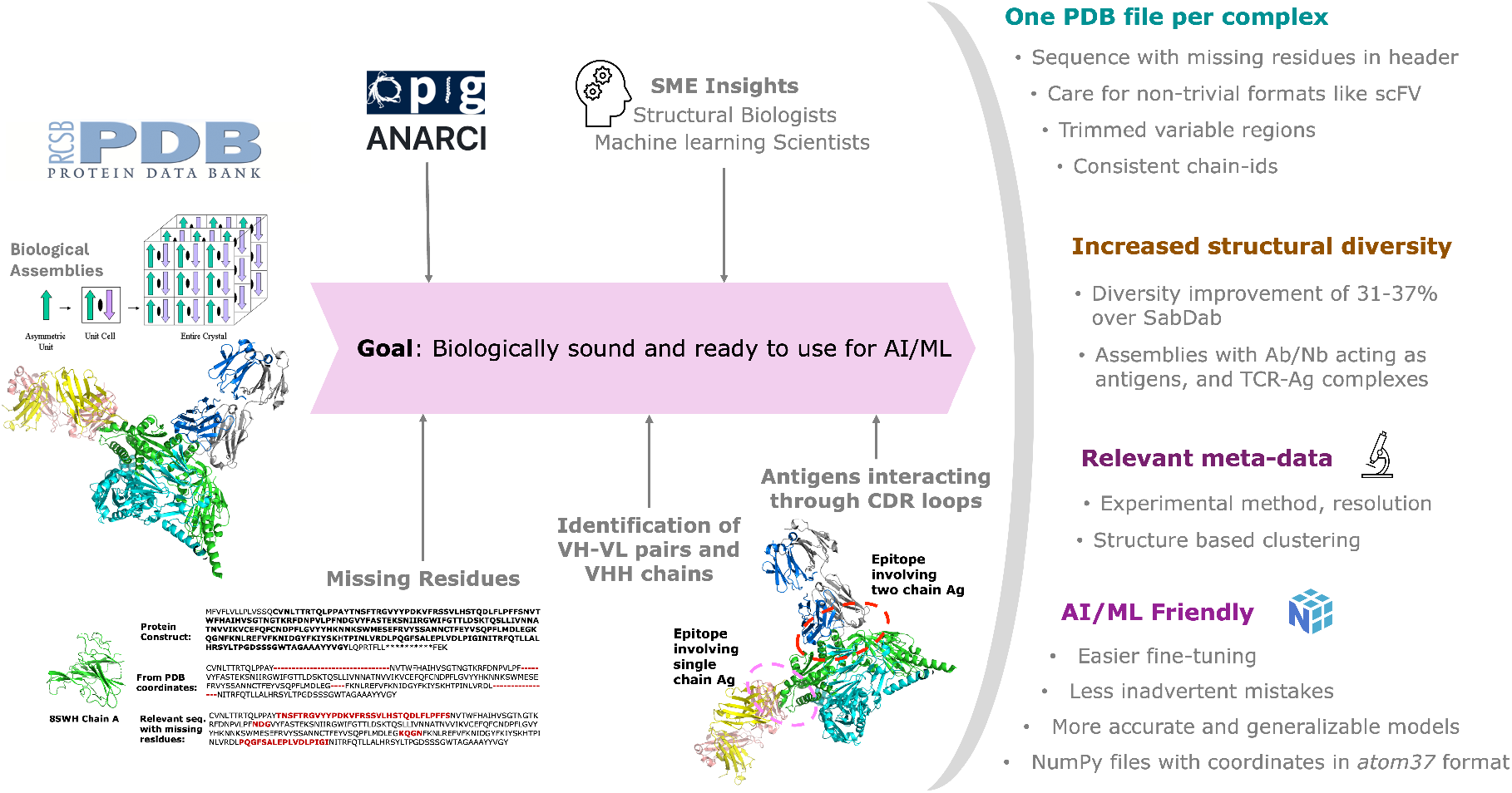
Overview of the SNAC-DB Data Curation Pipeline. Public PDB biological-assemblies are downloaded and parsed to recover missing residues, then annotated with IMGT numbering via ANARCI to define variable chains. A series of structural-biology–informed rules are applied to identify Ab–Ag and Nb–Ag complexes, and converted to ML-ready formats. The right panel highlights some key features of the curated data. The RCSB PDB logo is obtained from https://www.rcsb.org/, and the OPIG ANARCI logo from the Oxford Protein Informatics Group website https://opig.stats.ox.ac.uk/.

We demonstrate SNAC-DB’s impact through three key contributions. First, using a rigorous benchmark of all public PDB deposits from May 2024 to March 2025 plus confidential therapeutic complexes (Figure 2), we reveal that seven state-of-the-art (SOTA) models (Protenix-v1, OpenFold-3p2, RosettaFold-3, Boltz-2, Boltz-1x, Chai-1, and AlphaFold2.3-multimer) rarely exceed 25% success rates on novel targets, with systematic overestimation in published benchmarks. Second, we quantify a critical gap between model sampling capability and confidence-based ranking: while generating 1000 samples per target substantially increases the likelihood of producing accurate structures (oracle selection: 50.5%), confidence-based ranking remains nearly flat (10.9–14.9%), revealing that improved ranking mechanisms represent a more tractable path to performance gains than simply scaling sample counts. Third, we validate SNAC-DB’s practical value by fine-tuning GeoDock on our expanded dataset, achieving higher success rates compared to SAbDab training (11.0% vs. 7.1% for antibodies; 7.0% vs. 4.0% for NANOBODY^®^ VHHs). Together, these findings underscore both the limitations of current methods and the critical role of comprehensive, ML-ready data curation in advancing antibody–antigen structure prediction for therapeutic applications.

**Figure 2:**
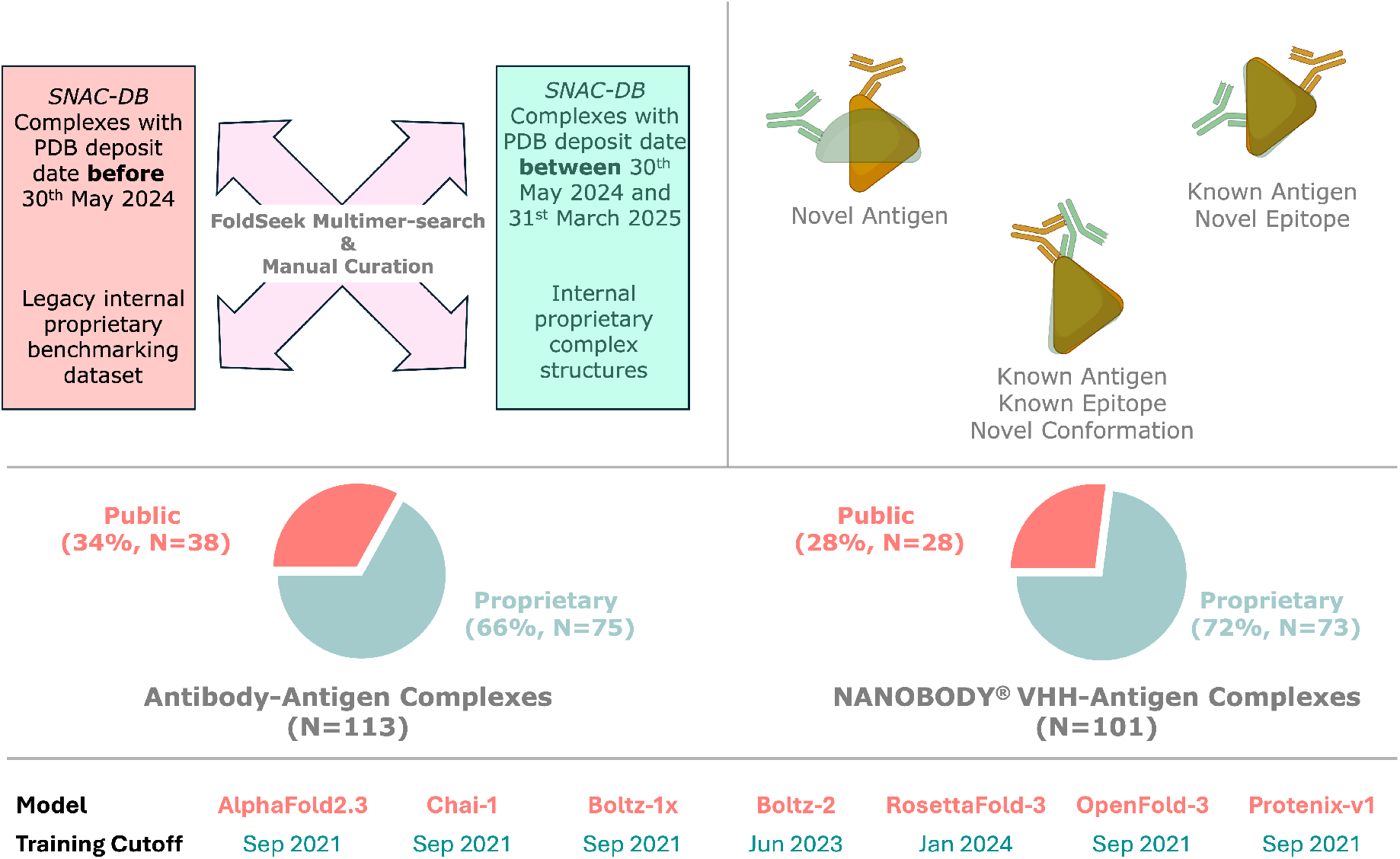
Overview of benchmarking dataset curation. *Top left:* The four input sources and filtering logic. SNAC-DB complexes deposited before May 30, 2024 and a legacy internal proprietary dataset are excluded via FoldSeek Multimer-search and manual curation to ensure no overlap with model training sets. The final benchmark draws exclusively from public PDB deposits between May 30, 2024 and March 31, 2025, and internal proprietary therapeutic structures. *Top right:* The three evaluation scenarios represented in the benchmark: novel antigens (antigen sequence unseen during training), known antigens with novel epitopes, and known antigens with known epitopes but novel binding conformations. *Mid left:* Composition of the antibody–antigen benchmark (N=113), comprising 34% public (N=38) and 66% proprietary (N=75) complexes. *Mid right:* Composition of the NANOBODY^®^ VHH–antigen benchmark (N=101), comprising 28% public (N=28) and 72% proprietary (N=73) complexes. *Bottom:* Training PDB cutoff dates for all evaluated models are listed at the bottom.

## 2 Results

As SAbDab is the *de facto* resource for curated Ab/Nb–Ag complexes, we directly compare its curation logic and dataset with those of SNAC-DB. We then evaluate the performance of SOTA complex-prediction models on our newly assembled benchmarking dataset.

### 2.1 Comparison with SAbDab

To demonstrate the advantages of SNAC-DB’s curation pipeline—such as improved capture of multichain antigens, a greater total number of complexes, and enhanced structural diversity—we perform a head-to-head comparison with SAbDab. Since SNAC-DB focuses exclusively on polypeptide antigens, we first preprocess the SAbDab download by removing any entries composed solely of non-protein ligands (e.g., haptens, carbohydrates, nucleic acids). For mixed-composition complexes, we retain only the polypeptide chains. We further restrict both datasets to PDB deposited before March 31, 2025.

#### 2.1.1 Pipeline Logic

To isolate differences in curation logic, we applied the SNAC-DB pipeline to the asymmetric units of the 8,670 PDB entries for which SAbDab reports at least one complex with a polypeptide antigen (excluding entries that failed during download or ANARCI processing and those containing unbound light chains). Both pipelines catalogued 16,527 complexes in total, including cases where no antigen was detected.

The first point of divergence occurs in 733 entries where SAbDab finds no antigen but our pipeline does—primarily capturing complexes in which antibodies or NANOBODY^®^ VHHs serve as antigens, such as Anti-Fab VHHs, VHH-Fabs, etc., as demonstrated in Figure 3a for PDB 8T60 (Hariharan et al., 2024). But more importantly, our pipeline identifies all antigen chains contributing to a single epitope more accurately. Capturing full epitope–paratope interfaces is essential, since biologically a complex would not form if any epitope chain were omitted. For example, PDB 7PA8 (Harprecht, 2020) is a homo-pentamer; Figure 3b contrasts how SAbDab and our pipeline handle it. SAbDab treats each antibody as binding a single subunit—reporting a one-chain antigen—whereas inspection shows the true epitope spans two adjacent subunits. Consequently, our method correctly assigns a two-chain antigen. Figure 4a compares the distribution of complexes by antigen chain count (0–5 chains). As expected, our pipeline identifies fewer unbound VH–VL/VHH chains and single-chain antigens, with most additional gains occurring in two-chain antigens, while even reduction in five-chain ones. These criteria strike a balance—recovering true multi-chain epitopes without overestimating chain membership. Capturing such nuances will help ML models learn the true drivers of complex formation and enhance traditional bioinformatics analyses.

**Figure 3:**
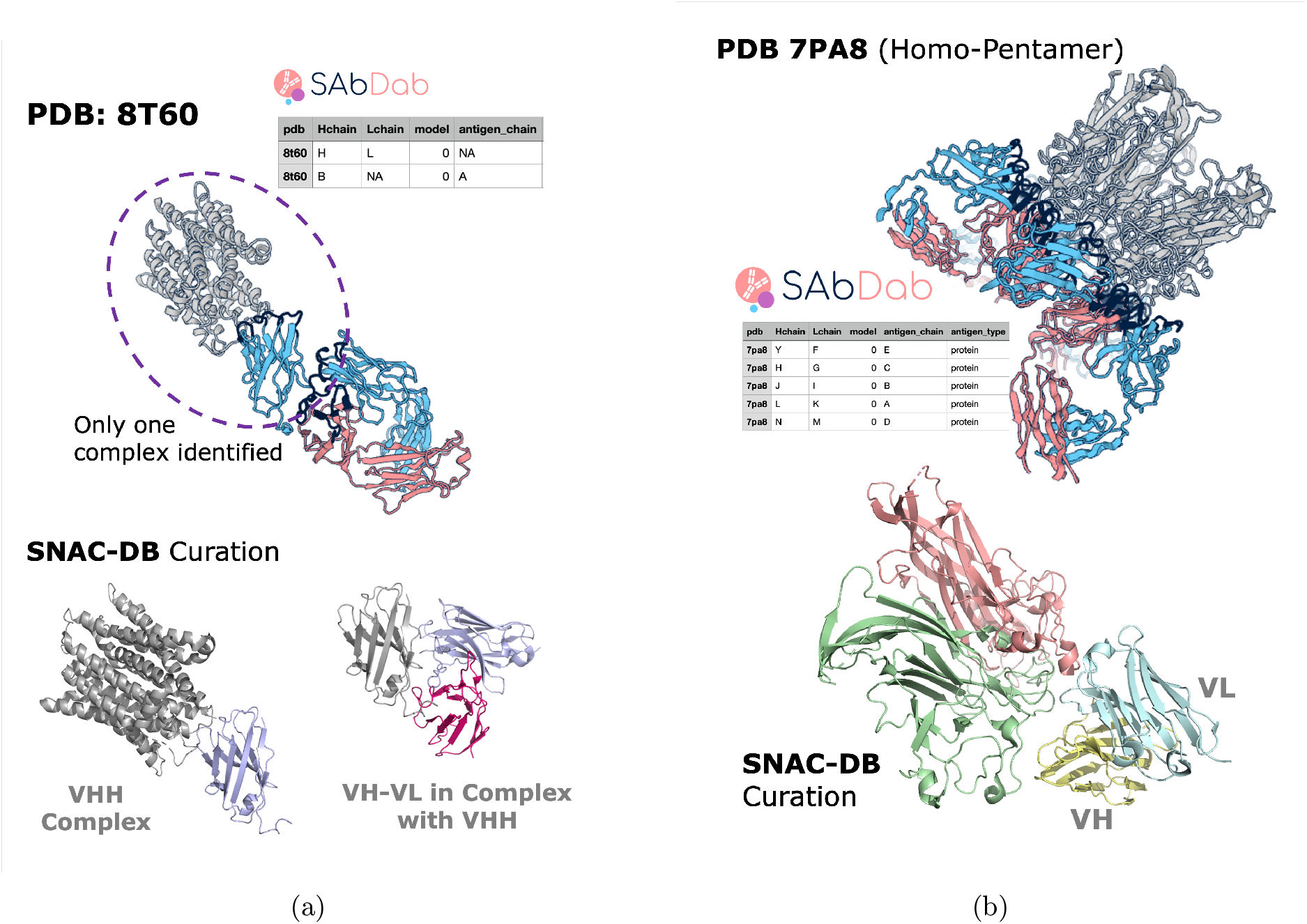
SNAC-DB Pipeline Captures Often-Missed Complexes and Multi-Chain Epitopes. (**a**) **Ab/Nb acting as antigens**. Top: SAbDab’s annotation of PDB 8T60, with VH/VHH chains in blue and VL chains in salmon. Top-right: Table of the two complexes SAbDab identifies—one of which is an unbound antibody. Bottom: SNAC-DB’s annotation of the same PDB, identifying a VHH complex, and a VH-VL bound to a VHH. (**b**) **Accuracy for Multi-chain Epitopes**. Top: SAbDab’s annotation of PDB 7PA8, with VH chains in blue and VL chains in salmon. Middle-left: Table of the five complexes SAbDab identifies—each pairing an antibody with a single antigen chain. Bottom: SNAC-DB’s annotation of the same PDB, showing the antigen epitope spanning two subunits of the homo-5-mer (pale green and wheat), correctly capturing the multi-chain interface.

**Figure 4:**
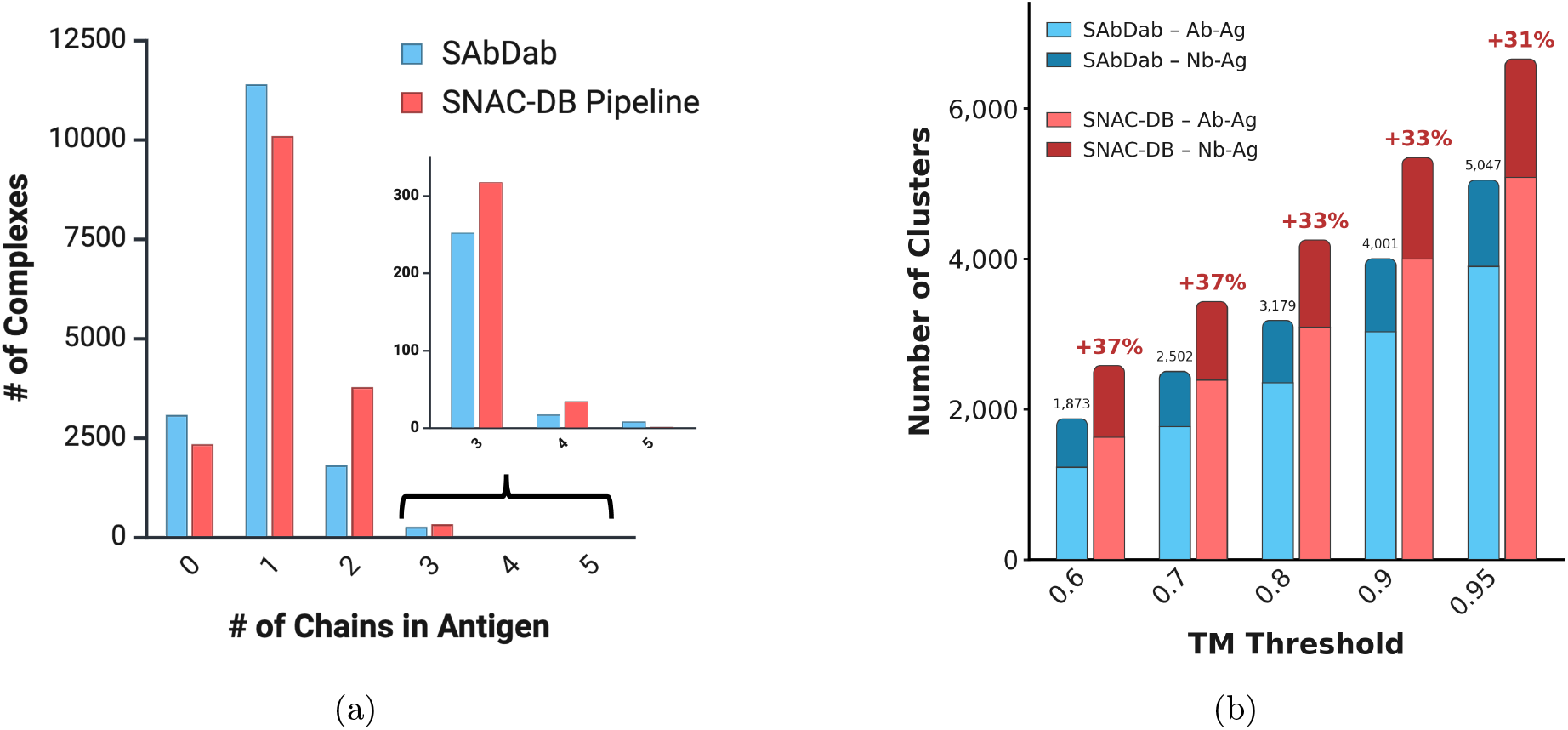
SNAC-DB Provides Greater Structural Diversity. (**a**) **Antigen Chain-Count Comparison**. Bar plot comparing the number of complexes with 0–5 antigen chains identified by SAbDab (blue) versus the SNAC-DB pipeline (red) across 16,527 curated entries (last PDB deposit date of March 31, 2025). A count of 0 indicates unbound antibody or NANOBODY^®^ VHH structures. An inset zooms in on the 3–5 chain counts for clarity. (**b**) **Structural Diversity Comparison**. Stacked bar plot comparing the number of structural clusters—computed via FoldSeek-Multimer at varying TM-score thresholds—for SAbDab (blue) and SNAC-DB (red), with clusters decomposed into Ab-Ag (light) and Nb-Ag (dark) based on majority composition of cluster members. Only complexes containing at least one protein antigen chain are included, and for SAbDab entries the Ab/Nb chains were pre-trimmed to their variable regions. Percentage increase in total cluster count for SNAC-DB relative to SAbDab is annotated above each bar pair. Both datasets are limited to PDB deposits up to March 31, 2025.

#### 2.1.2 Structural Diversity

Next, we compare the final SNAC-DB dataset against the bound complexes identified by SAbDab. Processing all biological-assemblies gives SNAC-DB more entries, 15,523 vs. 13,919 for SAbDab. For a fair comparison, we trim the Ab/Nb chains of the SAbDab identified complexes to retain only the variable regions, and apply Foldseek-Multimer clustering at multiple TM-score similarity thresholds to both datasets, counting the resulting clusters. As shown in Figure 4b, SNAC-DB preserves significantly greater structural diversity than SAbDab which is consistently maintained above 30% across TM threshold, indicating its potential to substantially augment the training and fine-tuning data for current SOTA methods by up to 37% more relevant structures.

While acquiring new experimentally determined Ab–Ag and Nb–Ag structures remains crucial for improving prediction accuracy and advancing biologics discovery, such efforts are costly and time-consuming. By offering an immediately available dataset with enhanced structural coverage, SNAC-DB can accelerate model development and deliver impact without waiting for future structure deposits.

### 2.2 Benchmarking SOTA Models

We evaluated seven recently released ML models that have raised hopes for significantly improved Ab–Ag and Nb–Ag complex prediction: Protenix-v1 (Team et al., 2026), OpenFold-3p2 (OF3p2; (The OpenFold3 Team, 2025a,b)), RosettaFold-3 (RF3; (Corley et al., 2025)), Boltz-2 (Passaro et al., 2025), Boltz-1x (Wohlwend et al., 2024), Chai-1 (Chai Discovery, 2024), and AlphaFold2.3-multimer (Evans et al., 2021). We also intended to include AlphaFold3 (Abramson et al., 2024), but licensing restrictions precluded its direct use. Given its architectural kinship, we hope that the combined performance of Protenix-v1, OF3p2, RF3, Boltz-2, Boltz-1x, Chai-1 should be representative of AlphaFold3’s accuracy. All predictions used default parameters without additional restraints or auxiliary information.

Figure 2 presents an overview of our benchmarking dataset curation workflow alongside the training cutoff dates of the seven models evaluated. We assess prediction quality using DockQ (Mirabello and Wallner, 2024), the standard metric for benchmarking ML-predicted protein–protein complex quality against experimentally determined structures. Accurate evaluation of antibody–antigen predictions requires careful handling of multi-chain epitopes, where the binding interface may span multiple antigen subunits; naively applying DockQ to such cases risks over- or underestimation of performance. We developed a custom methodology to address this, detailed in Supplementary Section S4. DockQ scores are categorised as Incorrect (*<*0.23), Acceptable (0.23–0.49), Medium (0.49–0.80), and High (≥0.80). Our benchmark draws exclusively from structures deposited after May 2024, ensuring no overlap with any model’s documented training data (*bottom* of Figure 2), though undisclosed validation sets used during model development may overlap with our benchmark.

#### Ab–Ag prediction performance

On the full antibody–antigen benchmark (Figure 5a), Protenix achieved the highest success rate at 23.0%, followed by RF3 at 21.2%, with Boltz-2, AlphaFold2.3, OF3p2, Boltz-1x, and Chai-1 trailing at 19.5%, 16.8%, 16.8%, 14.2%, and 13.3%, respectively. All models except Boltz-1x produced some high-quality predictions (DockQ ≥ 0.80), ranging from 0.9% to 5.3%. When separating public from proprietary complexes, Chai-1 and Boltz-1x perform notably better on proprietary structures than on public ones, while the gap narrows for other models and reverses for Protenix. To contextualise this, Figure S7 shows the query-normalised TM-score of each benchmark complex against its nearest neighbour in the SNAC-DB training set (defined here as all SNAC-DB complexes deposited before May 30, 2024), providing a measure of structural novelty relative to what models may have encountered during training. Despite Ab–Ag public and proprietary complexes showing broadly similar median TM-scores, the public set is considerably smaller (n=38 vs. n=75), meaning that its low-TM outliers represent a proportionally larger fraction of structurally challenging cases. Directly comparing these models is difficult given differences in architecture, training data, and training strategy, making it hard to attribute improvements over AlphaFold2.3 to any single factor; notably, only 3 of the 6 AlphaFold3-style models outperform AlphaFold2.3. Performance differences may also be partly driven by antigen class, with some models handling specific target types more effectively than others, such as membrane-bound antigens (Figure 7), though properly quantifying such effects would require balanced representation of major antigen categories in the benchmark, which is difficult to achieve in practice.

**Figure 5:**
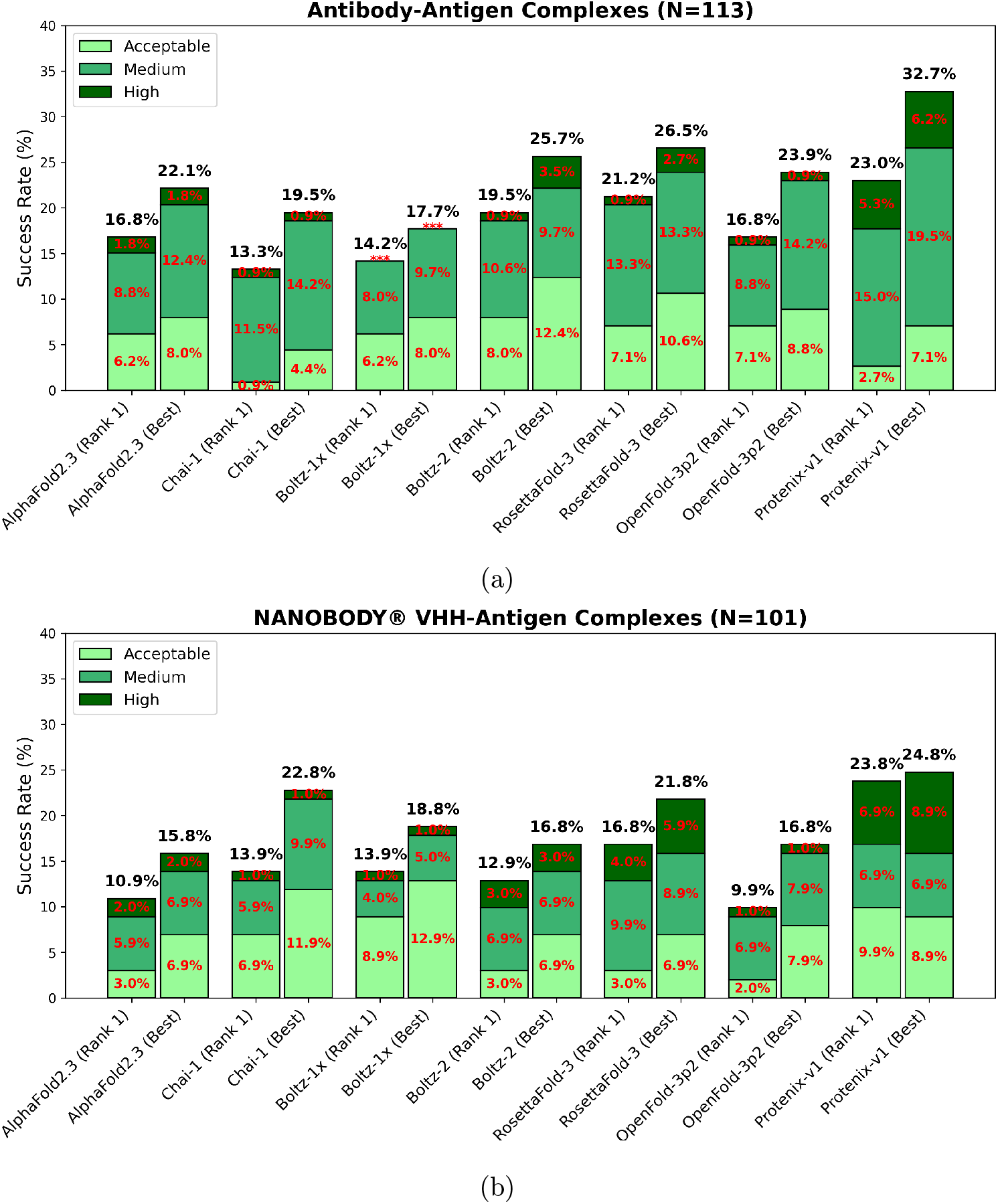
SOTA Performance. The bar plots display DockQ-classified performance (Acceptable (0.23 ≤ DockQ *<* 0.49), Medium (0.49 ≤ DockQ *<* 0.80), and High (DockQ ≥ 0.80)) of seven SOTA models (Protenix-v1, RosettaFold-3 (RF3), OpenFold-3p2 (OF3p2), Boltz-2, Boltz-1x, Chai-1, and AlphaFold ≥ 2.3-multimer) on **(a)** Ab–Ag and **(b)** Nb–Ag complexes, with the total number of samples mentioned in the respective titles. At the top of each bar, the total percentage of complexes with DockQ ≥ 0.23 is annotated, along with the respective breakdown for each of the DockQ categories. Categories with zero complexes are marked by **\*\*\***. Bars annotated “(Rank 1)” correspond to the top-ranked prediction by each model’s internal confidence metrics, while “(Best)” denotes the highest-quality prediction among five samples.

#### Nb–Ag prediction performance

On NANOBODY^®^ VHH–antigen complexes (Figure 5b), Protenix achieved the highest “Rank 1” success rate at 23.8%, with a comfortable margin over the other models, while OF3p2 and AlphaFold2.3 recorded the lowest at 9.9%. This margin shrinks considerably when comparing oracle-selected “Best” samples, indicating that other models are capable of sampling correct binding poses but fail to rank them highly. When split between public and proprietary complexes (Figure 6b), all models except AlphaFold2.3 perform significantly better on public complexes than on proprietary ones, suggesting that gains observed on public benchmarks do not consistently transfer to therapeutically relevant targets. This is not straightforwardly explained by structural novelty alone, as public and proprietary complexes show broadly similar median TM-scores relative to the SNAC-DB training set (Figure S7). A systematic difference we identified is a higher fraction of VHHs with longer CDR-H3 loops in proprietary complexes compared to public ones, illustrating the gap between publicly available structures and therapeutically relevant scenarios for this emerging modality.

**Figure 6:**
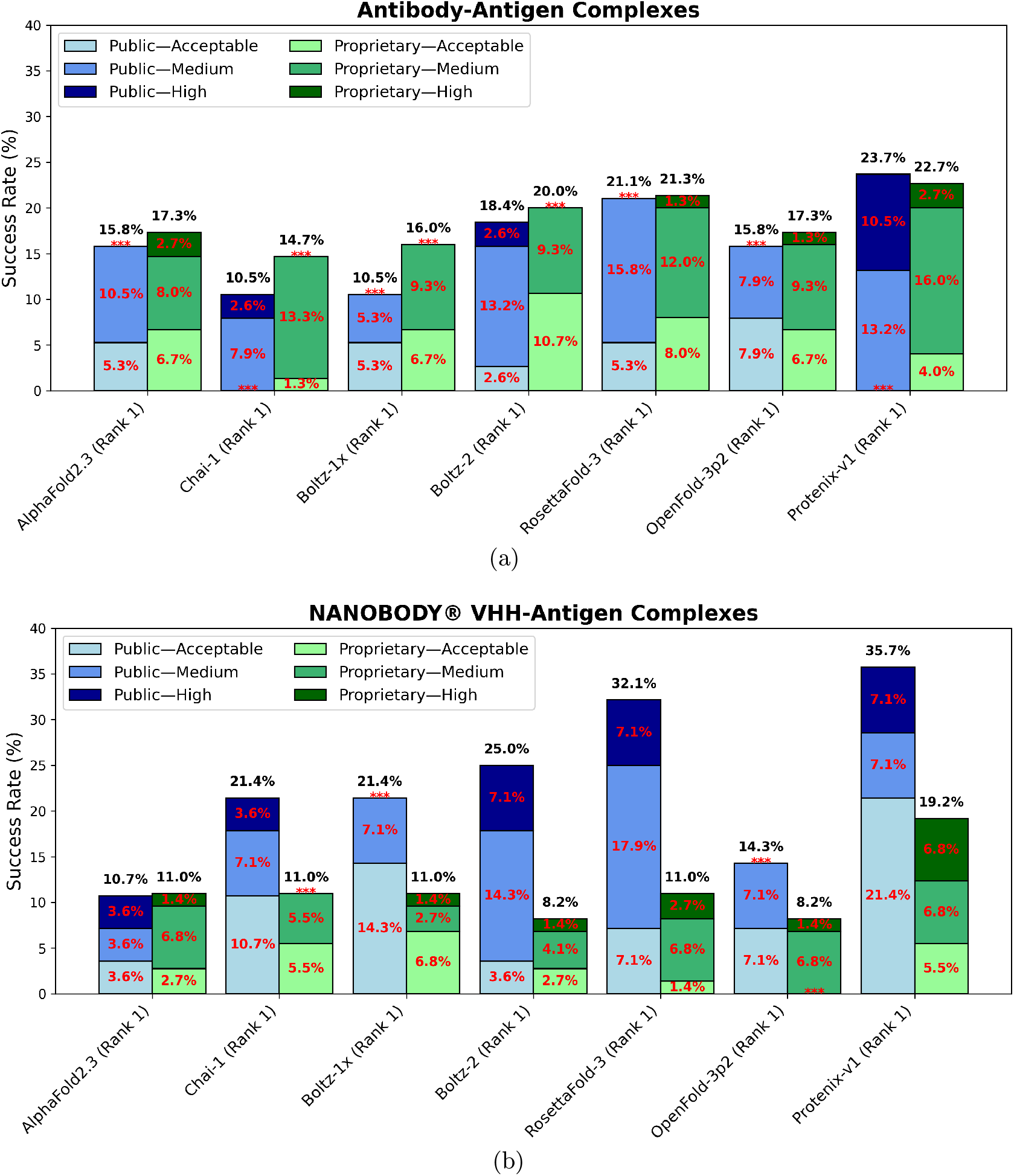
SOTA Performance on Public & Proprietary Benchmarking Complexes Separately. Stacked bar plots showing DockQ-classified “Rank 1” success rates for seven models, separated by public PDB (shades of *blue*), proprietary (shades of *green*) complexes. **(a)** Ab–Ag complex performance. **(b)** Nb–Ag complex performance.

**Figure 7:**
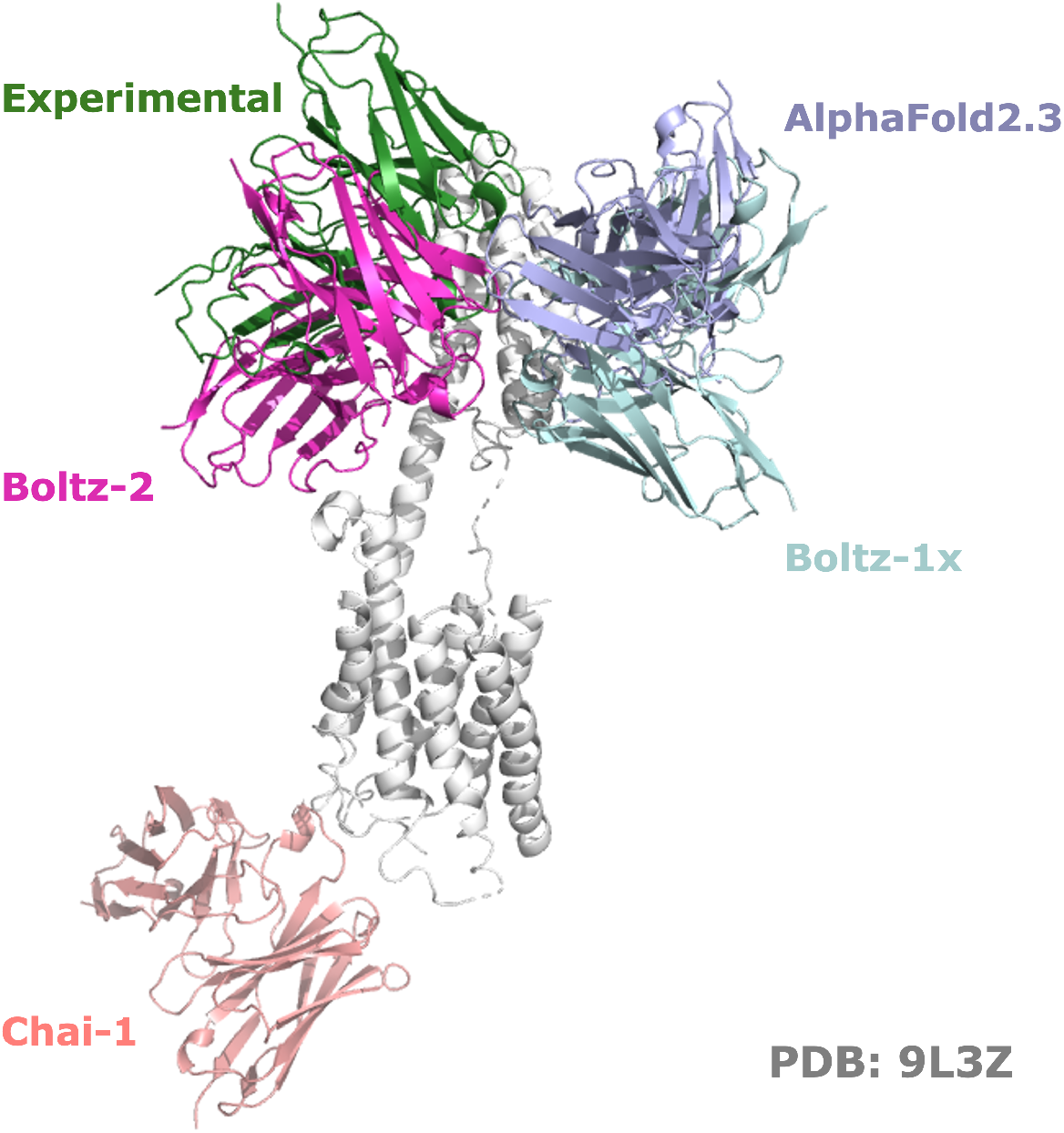
Performance on a membrane-bound antigen (PDB 9L3Z; (Lin et al., 2025)). Comparison of the experimental 9L3Z structure with model predictions: only Boltz-2 acceptably recapitulates the CDR–antigen interface, whereas all others misposition the binding site, illustrating divergent performance on challenging membrane-bound antigens.

#### Generalization performance

Figure 8 shows each model’s performance across the three test scenarios defined in Section 4.2. To enable comparison against documented training data, we computed the TM-score of each benchmark complex against its nearest neighbour in the SNAC-DB training set, classifying complexes as novel targets, known antigens with novel epitopes, or known antigens and epitopes with novel conformations. Performance varies across categories and models, with no single clear pattern, though one consistent trend emerges: novel antigen complexes yield almost no high-quality predictions across all models, whereas complexes involving familiar antigens (novel epitope or novel conformation) achieve substantially higher success rates. This is consistent with models relying heavily on learned structural priors for the antigen: when the antigen fold is represented in training data, the model can leverage that knowledge regardless of whether the epitope location or bound conformation is new, whereas an entirely unseen antigen fold removes this scaffold and the binding interface must be predicted with little structural guidance. Overall, Protenix shows the most balanced performance across all three scenarios, making it the most reliable model in this evaluation. We note that training data for several models remain partially undisclosed, precluding definitive conclusions about generalization capabilities.

**Figure 8:**
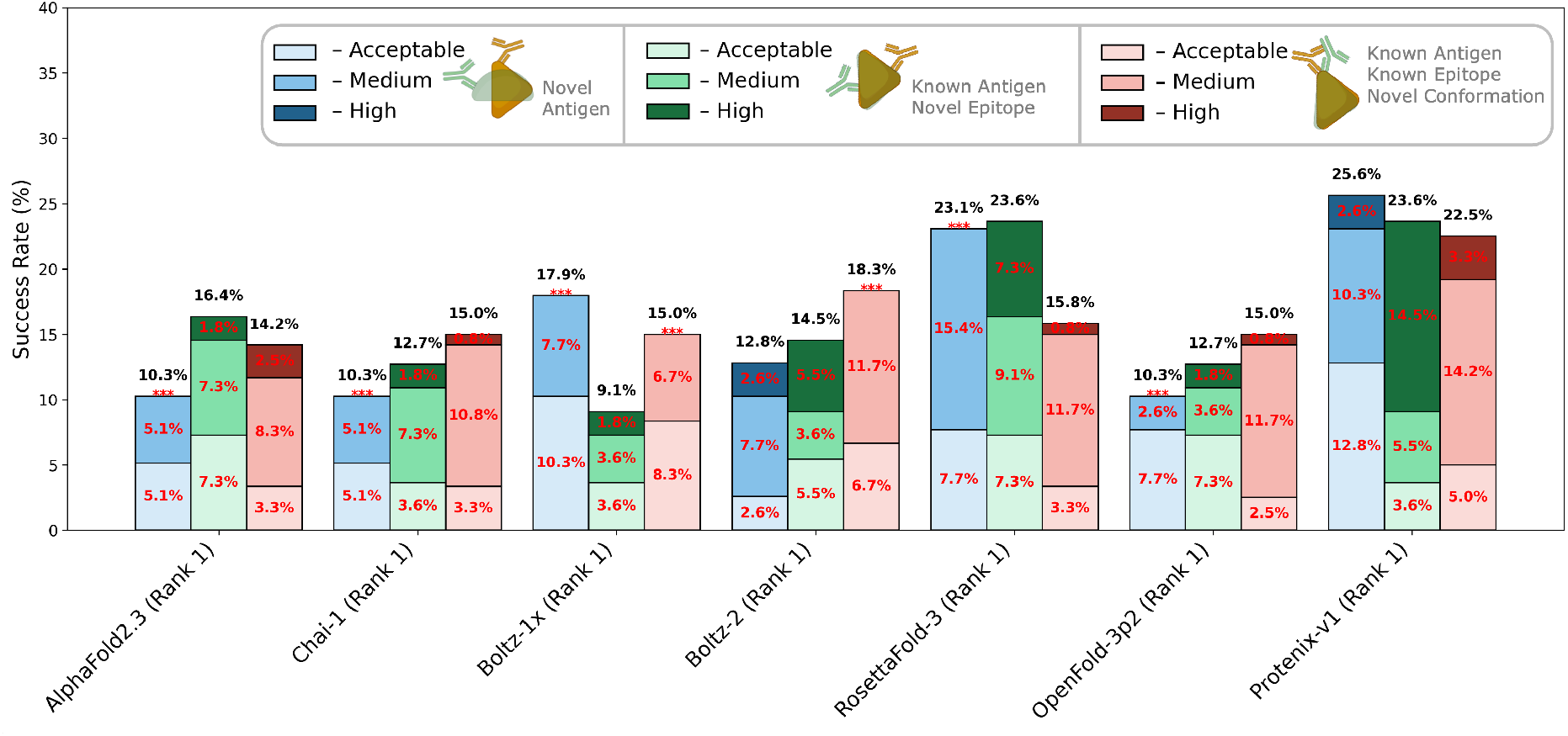
SOTA Performance on Different Test Scenarios. Stacked bar plots of DockQ-classified “Rank 1” success rates (Acceptable, Medium, High) for combined Ab–Ag and Nb–Ag benchmarks, comparing the seven models. Bars are colored by test scenario—*blue*: **novel target**, *green*: known antigen with **novel epitope**, *pink* : known antigen and epitope with **novel conformation**. Scenarios were defined by comparing benchmark complexes against SNAC-DB PDB entries deposited before May 30, 2024.

#### Inbuilt confidence scores and ranking

Although AlphaFold–style models report internal confidence metrics (e.g., pLDDT) and generate ranking scores, they frequently miss the most accurate structure. In every case, the “best of five” success rate outperforms the top-ranked prediction—for instance, Boltz-2 achieves 25.7% success when evaluating its top five samples versus 19.5% for its highest-scoring model on the Ab–Ag set. The same pattern holds for all methods and for Nb–Ag complexes. These results suggest that, while the correct pose is often generated, the internal scoring functions do not reliably distinguish it as the top prediction.

To assess potential *overconfidence*, Figure 9a plots both the model’s internal confidence score and its interface predicted template modeling score (ipTM) against DockQ for Boltz-2’s top-ranked predictions. While low confidence or ipTM reliably flag inaccurate models, high values do not correlate with correctness (Spearman *ρ* = 0.41 for confidence and *ρ* = 0.36 for ipTM).

**Figure 9:**
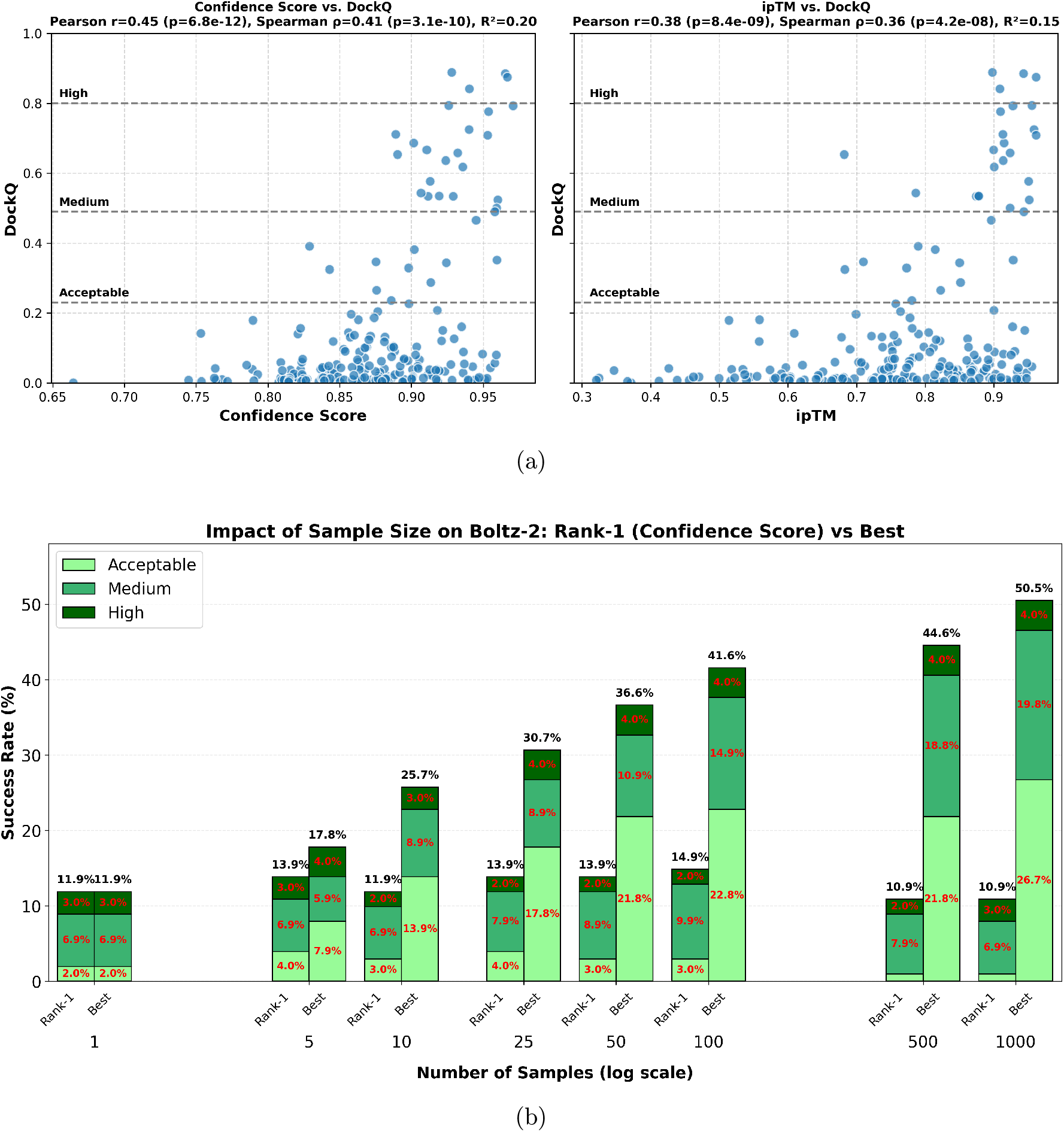
Confidence scoring limitations in Boltz-2 structure prediction. (a) **Confidence metrics vs. DockQ**. Scatter plots of top-ranked Boltz-2 predictions showing internal confidence score (left) and ipTM (right) against DockQ. Horizontal dashed lines mark DockQ thresholds: Acceptable (0.23), Medium (0.49), High (0.80). While low confidence reliably flags incorrect predictions, high confidence does not correlate with accuracy (Spearman *ρ* = 0.41 for confidence, *ρ* = 0.36 for ipTM). (b) **Sampling strategy impact**. Evaluation on NANOBODY^®^ VHH-antigen complexes (N=101) using 1-1000 samples per case comparing **Rank-1** (selecting the prediction with highest confidence score, as would be done in practice) versus **Best** (oracle selection of the prediction with highest DockQ score). While increased sampling improves the likelihood of generating accurate structures, confidence-based ranking fails to identify them: at 1000 samples, Rank-1 achieves only 10.9% success versus 50.5% for oracle selection, demonstrating that improved ranking mechanisms represent a more tractable path to performance gains than scaling sample counts.

#### Trust in real-world drug discovery

A recurring theme across our evaluation is the gap between performance figures reported in model publications and what we observe on an out-of-sample, therapeutically relevant benchmark. Published success rates of 40–60% on standard antibody– antigen test sets contrast sharply with the rates rarely exceeding 25% we find here, and the gap is more pronounced for NANOBODY^®^ VHH–antigen complexes, where all models achieve *<*20% success on proprietary structures, likely reflecting the greater structural complexity of therapeutically relevant NANOBODY^®^ VHH targets such as those with longer CDR-H3 loops that are underrepresented in current training data. This discrepancy is compounded by the unreliability of confidence scores: high confidence does not imply correctness, and practitioners cannot use model certainty as a reliable proxy for prediction quality. Together, these findings highlight that published benchmarks may paint an overly optimistic picture of deployment readiness, and that rigorous out-of-sample evaluation on proprietary, therapeutically relevant structures is essential before these methods can be routinely trusted in drug-discovery pipelines.

### 2.3 Sample Size Impact on Prediction Success

Since the publication of AlphaFold3 (Abramson et al., 2024), a critical question has emerged: can the accuracy of generative structure prediction models be improved simply by generating more samples per target? To address this, we systematically evaluated the impact of sample size on Boltz-2 performance using our NANOBODY^®^ VHH-antigen complex benchmark (N=101), generating 1, 5, 10, 25, 50, 100, 500, and 1000 samples per case. Predictions were generated in batches of 5 for configurations up to 100 samples and in batches of 25 for the 500 and 1000 sample scenarios, with each batch using a different random seed (while everything else identical) to ensure diversity in initialization.

As we did earlier, for each sample size, we compared two selection strategies (Figure 9b): (1) **Rank-1**, where we select the prediction with the highest model-reported confidence score, representing practical usage without access to ground truth; and (2) **Best**, where we oracle-select the prediction with the highest DockQ score, representing the theoretical upper bound achievable with perfect ranking.

The results reveal two critical insights. First, while increasing sample size substantially improves the likelihood of generating accurate binding poses (Best selection improves from 11.9% to 50.5% as samples increase from 1 to 1000), Rank-1 performance remains nearly flat (fluctuating between 10.9% and 14.9%), as noted earlier, demonstrating that current confidence scoring fails to identify correct structures even when they exist in the ensemble. This striking gap—where oracle selection achieves 4.5× better performance than confidence-based ranking at 1000 samples—indicates that improving ranking mechanisms could provide immediate and substantial performance gains without additional computational cost. This ranking deficiency persists even when using ipTM, an interface-specific confidence score predicted by the model (Unsal et al., 2025), as shown in Figure S8.

However, the limited ceiling of oracle performance (50.5% at 1000 samples) reveals a complementary challenge: even with perfect ranking, the underlying generative model frequently fails to produce accurate structures. Scaling to 1000 samples requires substantial computational resources (approximately 2-3 hours per case on a A100 GPU) while yielding diminishing returns given the apparent saturation of the Best curve. These findings suggest a two-pronged path forward: (1) in the near term, developing better confidence calibration and ranking mechanisms represents the most practical route to improved performance; (2) in the longer term, fundamental improvements to the generative models themselves are needed to increase the quality of sampled structures. Until ranking mechanisms are substantially improved, however, we see limited practical benefit in simply scaling up sample counts beyond current levels.

### 2.4 Impact of SNAC-DB on Model Performance

To demonstrate the practical value of SNAC-DB’s expanded structural diversity and improved curation, we fine-tuned GeoDock, a representative flexible docking ML model, using both SNAC-DB and SAbDab training sets. This experiment directly tests whether SNAC-DB’s enhanced coverage and principled sample weighting translate into measurable improvements in prediction accuracy.

#### Fine-tuning protocol

We adapted the GeoDock codebase to incorporate per-complex sample weights during training. For SNAC-DB, we applied the cluster-based weights provided with the database (inverse square root of cluster size at TM-score threshold 0.99). For SAbDab, we followed the identical clustering and weighting protocol used in SNAC-DB curation to ensure a fair comparison. Both models were initialized from the publicly available `dips_0.3.ckpt` checkpoint and trained for up to 200 epochs with early stopping. Training was performed on a single AWS node with 4 NVIDIA A10G GPUs (24GiB each), using the original GeoDock hyperparameters (learning rate 1e-5, batch size 1 per GPU). Early stopping selected the SAbDab fine-tuned model at epoch 116 and the SNAC-DB model at epoch 163, consistent with the larger and more diverse SNAC-DB training set requiring more epochs to converge.

#### Evaluation on benchmarking dataset

We evaluated both fine-tuned models on our rigorously curated benchmark dataset (Section 4.2), which was explicitly filtered to exclude structural overlap with both training sets. Since GeoDock requires separate structures for both ligand and receptor, we used Boltz-2 predicted apo Ab and Nb structures as input to ensure no information leakage from the bound complex. Figure 10a shows the performance breakdown by DockQ categories for antibody (Ab) and NANOBODY^®^ VHH (Nb) complexes. The SNAC-DB fine-tuned model achieved a 11.0% success rate (DockQ ≥ 0.23) on Ab–Ag complexes, compared to 7.1% for the SAbDab-trained model. For Nb–Ag complexes, SNAC-DB fine-tuning yielded 7.0% success versus 4.0% for SAbDab. As expected, both models substantially outperformed the original GeoDock (0% success on both complex types), confirming that task-specific fine-tuning on antibody and NANOBODY^®^ VHH data is essential for this application. More importantly, SNAC-DB’s consistently superior performance over SAbDab fine-tuning—achieving 1.5× higher success for Ab–Ag and 1.75× higher for Nb–Ag—demonstrates that training on diverse, structurally representative data enables more accurate modeling of biologically relevant binding geometries.

**Figure 10:**
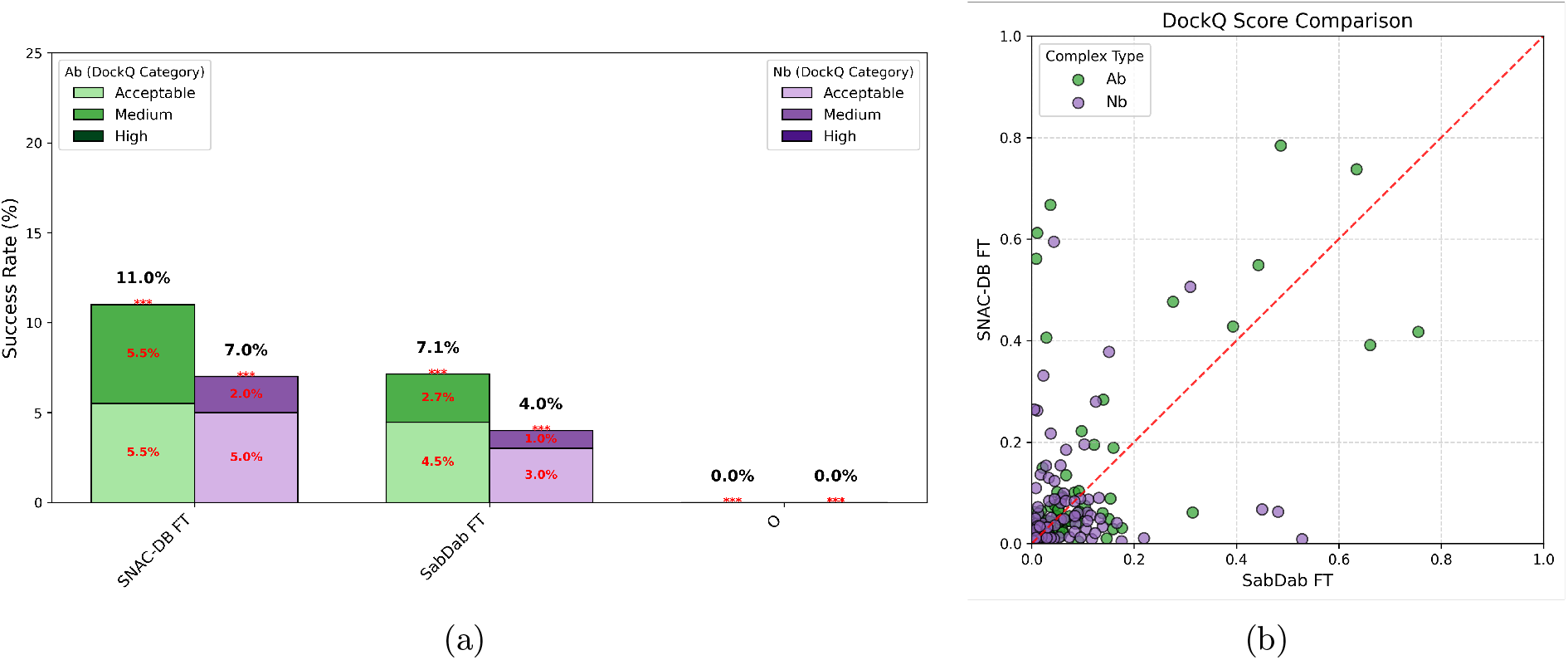
Impact of SNAC-DB on GeoDock Performance. (**a**) Stacked bar plots showing DockQ-classified performance of GeoDock models fine-tuned on SNAC-DB (left) versus SAbDab (middle) on our benchmarking dataset, separately for Ab–Ag (green) and Nb–Ag (purple) complexes. The right panel shows the original untrained GeoDock performance for reference. At the top of each bar, the total percentage of complexes with DockQ ≥ 0.23 is annotated, along with the respective breakdown for Acceptable (0.23 ≤ DockQ *<* 0.49), Medium (0.49 ≤ DockQ *<* 0.80), and High (DockQ ≥ 0.80) categories. (**b**) Head-to-head comparison of DockQ scores for each benchmark complex, with SNAC-DB fine-tuned model on the y-axis and SAbDab fine-tuned model on the x-axis. Points above the diagonal (red dashed line) indicate superior performance from SNAC-DB training. Green circles denote Ab–Ag complexes; purple circles denote Nb–Ag complexes. The majority of points lie above the diagonal, confirming consistent improvements from SNAC-DB’s expanded structural diversity and principled sample weighting.

Figure 10b presents a scatter plot comparing DockQ scores for each benchmark complex predicted by the two models. Points above the diagonal indicate cases where SNAC-DB fine-tuning outperformed SAbDab. The majority of points lie above the diagonal, confirming the consistent improvement from SNAC-DB training. The small number of cases where SAbDab performs slightly better likely reflect stochastic training effects and differences in how sample weighting affects learning dynamics, rather than fundamental gaps in data coverage.

#### Implications for model development

These results validate SNAC-DB’s core premise: expanded structural diversity and transparent redundancy control directly improve model generalization. The near doubling of rates for both Ab-Ag and Nb-Ag complexes—achieved without architectural changes or hyperparameter tuning—underscores the value of comprehensive data curation.

## 3 Discussion

SNAC-DB provides a vital resource for both structural biologists and the ML community to address the antibody and NANOBODY^®^ VHH–antigen complex prediction challenge—a persistent blind spot that limits the real-world impact of SOTA methods in drug discovery.

At its core, SNAC-DB is a comprehensive database of Ab–Ag and Nb–Ag complexes that fills critical gaps in existing resources and prevents researchers from discarding experimentally determined structures carrying valuable binding information. Beyond standard antigen–binder pairs, we include complexes where antibodies or NANOBODY^®^ VHHs serve as antigens, as well as TCR–pMHC assemblies typically overlooked by immunoglobulin-centric pipelines. Our methodology accurately recognizes multi-chain epitopes and retains complexes with weak CDR-loop crystal-packing contacts. By operating on author-defined biological-assemblies rather than asymmetric units, SNAC-DB ensures an immediate head start on identifying biologically relevant complexes. These features collectively enrich training and fine-tuning datasets, empowering ML models to learn the true determinants of immunoglobulin-mediated binding.

To streamline adoption, SNAC-DB is fully ML-ready: each complex is delivered as a cleaned PDB file and Python dictionary, with full sequences—including missing residues recovered via header–coordinate alignment—trimmed to variable domains and packaged as *atom37* NumPy arrays alongside unified metadata CSV files, eliminating common preprocessing pitfalls. We supply Foldseek-Multimer clustering assignments and cluster sizes at multiple TM-score thresholds, enabling principled sample weighting so that every structure can be leveraged during training.

A contact-level analysis of the interfaces reveals what underlies this structural diversity (Figure S5). On top of the gain in structurally distinct complexes demonstrated in Figure 4b, SNAC-DB contains more than twice the proportion of multi-chain antigen complexes as SAbDab (33.5% vs. 15.3%), reflecting explicit enumeration of all antigen chains within contact distance of the paratope. Epitope secondary-structure composition is more balanced, with a modestly lower loop fraction and proportionally more helix and strand contacts, and paratope CDR usage is slightly less H3-dominated across both antibodies and VHHs. At the sequence level, 2–10% of SNAC-DB complexes carry an antigen chain with no close SAbDab homologue (Figure S6), representing targets for which SAbDab provides no training signal. A detailed interface composition analysis is provided in Supplementary Section S2.3.

We rigorously evaluated seven cutting-edge ML models (Protenix-v1, RosettaFold-3, OpenFold-3p2, Boltz-2, Boltz-1x, Chai-1, and AlphaFold2.3-multimer) using a benchmark comprising all public PDB deposits from May 2024 to March 2025 plus confidential therapeutic structures. While success rates of 40–60% are often reported on standard datasets, our out-of-sample evaluation reveals systematic overestimation: even advanced methods struggle with novel targets and binding poses, rarely exceeding 25% success, and frequently misrank their best predictions. Obtaining reliable numbers required evaluation methodology tailored to antibody/NANOBODY^®^ VHH–antigen complexes, as standard DockQ applied naively risks averaging over irrelevant chain interfaces and masking incorrect binding modes at multi-chain epitopes, where per-chain scores can appear acceptable even when the predicted pose spans only a subset of the true interface (Section 4.3). The performance gap is particularly pronounced for NANOBODY^®^ VHH–antigen complexes, where proprietary therapeutic structures systematically underperform their public counterparts despite comparable structural novelty relative to the training set. Apart from the structural differences noted in the results, publicly deposited NANOBODY^®^ VHH structures are predominantly tool proteins used for conformational stabilization or as crystallography aids, with limited therapeutic relevance, meaning that models trained on public data are optimized for a very different distribution of binding scenarios than those encountered in drug discovery. This underscores that the field does not only need more structural data, but fit-for-purpose data reflecting the characteristics of therapeutically relevant binders.

Our systematic analysis of sampling strategies reveals two fundamental challenges facing current generative models. First, while increasing samples from 1 to 1000 substantially improves the likelihood of generating accurate structures (oracle selection: 11.9% → 50.5%), confidence-based ranking remains nearly flat (between 10.9% and 14.9%). This striking gap, where perfect ranking achieves nearly 4.5 times better performance than confidence scoring at 1000 samples, indicates that improved ranking mechanisms represent a more tractable path to performance gains. Second, the limited ceiling of oracle performance (50.5% at 1000 samples), combined with the substantial computational cost of large sample sets, reveals that simply scaling sample counts is insufficient and that fundamental improvements to generative model architectures are needed to increase the intrinsic quality of sampled structures. We also note that our benchmark is intentionally enriched for challenging, therapeutically relevant targets, and some failures likely reflect inherent difficulty in the experimental data themselves, including limited resolution, conformational disorder, and alternative binding modes, rather than purely computational shortcomings.

We validate SNAC-DB’s practical impact by fine-tuning GeoDock on our expanded dataset, achieving higher success rates compared to SAbDab training (11.0% vs. 7.1% for antibodies; 7.0% vs. 4.0% for NANOBODY^®^ VHHs). This demonstrates that SNAC-DB’s enhanced structural diversity directly translates to improved model generalization without architectural changes or hyperparameter tuning, confirming the value of comprehensive data curation for advancing prediction accuracy.

SNAC-DB and its publicly disclosed benchmark subset are fully reproducible and freely available, offering a one-stop resource for model development, evaluation, and comparison. By releasing all non-proprietary complexes from our benchmarking dataset alongside the ML-ready SNAC-DB with 37% expanded structural diversity, we provide both immediate practical improvements and a clear roadmap for future advances. In the near term, improved confidence calibration and ranking mechanisms can unlock substantial performance gains from existing models. In the longer term, enhanced generative architectures trained on diverse, high-quality data like SNAC-DB will be essential for achieving the accuracy and reliability required for routine deployment in therapeutic antibody discovery pipelines.

## 4 Materials and Methods

In this section, we first present our data curation pipeline—from downloading PDB entries via the RCSB PDB (Burley et al., 2025) to identifying and extracting individual complexes—detailing the rationale for each step. We then describe the filtering criteria and methodology used to assemble our benchmarking dataset.

### 4.1 Data Curation Pipeline

Each PDB entry includes both an asymmetric unit (ASU)—the smallest crystallographic repeat as deposited—and one or more biological-assemblies, which represent the author-validated (or software-predicted) functional oligomeric states. Biological-assemblies may consist of a single ASU, multiple ASUs (e.g., viral capsids), or a subset of an ASU (RCSB Protein Data Bank, 2025). We download all biological-assemblies for each PDB ID released by December 31, 2025. Unlike resources such as SAbDab that default to ASUs, our use of biological-assemblies ensures we begin with author-curated, biologically relevant complexes—and when no distinct bio-assembly is provided, the result simply falls back to the ASU—giving us at minimum a partially curated dataset to work from.

#### 4.1.1 Initial cleanup and annotation

We parse every PDB file associated with bioassemblies (and ASUs for viral capsids), retaining only polypeptide chains and atoms corresponding to standard amino-acid residues. Because regions with insufficient electron density are omitted from the coordinate section, residue numbering often “jumps” where residues are missing. However, the complete chain sequence is provided in the header, corresponding to the expressed protein. For each chain, we therefore extract the full sequence from the header and align it to the sequence derived from the coordinates—introducing gaps where density is absent. This alignment recovers the most accurate sequence information for each chain; any remaining missing residues are labeled “UNK” (or “X”) in the parsed output. We additionally preserve all the other meta-data from the header and associated b-factor values for each atom.

We then apply the widely used bioinformatics tool ANARCI (Dunbar and Deane, 2016) to each chain, annotating variable-region segments for antibody heavy, kappa, and lambda chains, as well as TCR *α, β, δ*, and *γ* chains. Each identified variable region is copied, assigned a new standardized chain identifier (relative to the original chain ID), and annotated with the ANARCI-derived IMGT numbering (Manso et al., 2022). We focus on extracting only the variable region (VH, VL, and VHH), as it is the primary determinant of antigen recognition via the CDR loops.

#### 4.1.2 Identifying VH-VL pairs and VHH

A PDB entry may contain multiple heavy and light chains, requiring identification of correct VH–VL pairings. We examine contacts across structurally conserved regions involved in VH–VL interactions: CDR3, framework regions FR2 and FR3 (which directly participate in the VH–VL interface), and portions of FR1 and FR4 (included to reduce false positives from crystal artifacts or atypical orientations). Using C_*α*_–C_*α*_ distances, we compute contact maps between VH and VL residues in these regions at distance cutoffs of 8 Å, 10 Å, and 12 Å. VH–VL pairs are assigned when consistent spatial proximity is observed across multiple regions and thresholds. This multi-region, multi-threshold strategy ensures sensitivity to conformational variation while minimizing false positives from crystal packing or antibody replicates.

PDB entries containing only heavy chains, or with unpaired heavy chains remaining after VH–VL pairing, are classified as NANOBODY^®^ VHH variable regions (VHH).

#### 4.1.3 Antigen Assignment

Identifying correct antigen partners for each VH–VL pair or VHH is the most critical stage of the pipeline. Through iterative refinement guided by structural biologists with over a decade of experience, we developed heuristics that closely match expert manual curation. All interactions are defined by contacts occurring through the CDR loops.

Our antigen detection strategy consists of two sequential stages. In the **complex identification stage**, we apply a permissive 12 Å C_*α*_–C_*α*_ distance cutoff across all CDRs to maximize sensitivity. This generous threshold accounts for resolution variability, coordinate uncertainty, and crystal packing distortions, ensuring we capture all potential antigen chains including those with weak or peripheral contacts.

In the subsequent **complex refinement stage**, we apply stricter 8 Å validation criteria empirically optimized to distinguish genuine binding interfaces from crystal-packing artifacts. The specific rules depend on ligand type (antibody or NANOBODY^®^ VHH) and whether the antigen is singlechain or multi-chain. For antibody complexes with single-chain antigens, at least one CDR contact from either VH or VL is required. For multi-chain antigens, both VH and VL chains must contact the antigen complex. When one chain of a multi-chain antigen meets the 8 Å criteria, the other chains identified at 12 Å are retained to preserve epitope integrity. Exceptions accommodate cases where VH-VL pairs act as antigens or where only one chain of a VH-VL ligand directly contacts the antigen. For NANOBODY^®^ VHH complexes, single-chain antigens require at least one CDR contact, while multi-chain antigens require contact excluding CDR1.

We also retain weak crystal-packing contacts involving CDR loops and annotate them explicitly, allowing users to filter these entries based on their training objectives. Finally, if no valid antigens pass the 8 Å validation and additional chains are present, we perform a rescue pass applying the same rules at 12 Å to recover poorly resolved or atypical complexes.

#### 4.1.4 Outputs

Upon completion of cleaning and parsing, the pipeline generates three core outputs. First, it writes cleaned PDB files in which chain labels have been standardized—heavy chains (“VH” and “VHH”) are relabeled as “H”, light chains (“VL”) as “L”, and antigen chains are assigned identifiers from “A” onward (skipping “H” and “L”, with numeric “0–9” and lowercase “a–z” used if more than 24 antigen chains are present). Missing residues—regions with insufficient electron density omitted from the original coordinate section but specified in the header—are incorporated into the sequence with their identities marked where known or labeled as “X” (unknown) where ambiguous. Second, it produces a NumPy (“.npy”) archive containing a top-level dictionary with the keys “VH”, “VL”, and “Ag”. Each of these entries maps to a nested dictionary of chain IDs, and for each chain we store the full aminoacid sequence (including resolved missing residues), *atom37* -formatted atomic coordinates, B-factors, and (for “VH” and “VL”) ANARCI-derived IMGT numbering. Finally, a summary CSV consolidates metadata for every complex—including the experimental method, resolution, deposition and release dates, and the assigned “VH”, “VL”, and “Ag” chain IDs—facilitating straightforward filtering and downstream analysis.

To eliminate redundant structures arising from multiple biological assemblies or asymmetric units of the same PDB entry, we applied FoldSeek-multimersearch to identify near-identical structures (template-modeling (TM) score > 0.999) and retained only the highest-priority structure based on completeness (number of residues, missing C_*α*_ atoms) and biological assembly preference.

Additionally, to facilitate sample weighting during model training, we perform structural clustering using Foldseek-Multimer at multiple TM-score based similarity thresholds (0.6, 0.7, 0.8, 0.9, 0.95, 0.99). We capture each complex’s cluster assignment and cluster size in a CSV manifest and supply a representative structure for every cluster. *As a good starting point, we suggest weighting samples inversely to the square root of their cluster size*.

### 4.2 Benchmarking Dataset Curation

The benchmarking dataset was designed to evaluate SOTA models under real-world conditions relevant to biotherapeutics development. To avoid overlap with model training data, we selected all public PDB entries deposited between May 30, 2024 and March 31, 2025, and supplemented them with proprietary, experimentally determined structures. Rather than relying on sequence-identity thresholds to define novelty, we emphasize structural dissimilarity when selecting VH–VL/VHH-antigen complexes.

Drawing on industry experience, we identified three key test scenarios:

1. **Novel targets**: Antigens for which no bound Ab/Nb structures existed in the training sets.
2. **Novel epitopes**: Targets present in the training data, but with previously unseen epitopes, to verify that models do not simply reproduce learned binding sites.
3. **Novel conformations**: Complexes with the same antigen and partially overlapping epitope as training examples, but featuring variations in Ab/Nb conformation, to assess the models’ ability to detect more fine-grained structural differences.

We maintained an approximate 20 / 25 / 55 split across these categories to ensure balanced representation and rigorous evaluation.

In addition to public PDB entries deposited before May 30, 2024, we included a legacy proprietary collection—together forming the *reference set*, against which we filtered our benchmarking set. The curation workflow consisted of four sequential stages using Foldseek-Multimer. **Stage 1 (Novel Targets):** We compared query antigens against reference antigens. Complexes with antigen TM-scores *<* 0.6 passed as novel targets. **Stage 2 (Novel Epitopes):** Failed complexes from Stage 1 underwent whole-complex comparison. Complexes with whole-complex TM-scores *<* 0.6 passed as novel epitopes. **Stage 3 (Novel Conformations):** Failed complexes from Stage 2 were compared using multi-chain structure alignment. Complexes with TM-scores *<* 0.85 passed as novel conformations. **Stage 4 (Redundancy Removal):** All passed complexes were compared against each other, retaining only structures with TM-scores *<* 0.85 to eliminate internal redundancy. All shortlisted structures underwent manual review to remove those with unresolved paratope/epitope coordinates or apparent crystal-packing artifacts. The final breakdown of benchmarking dataset complexes—from the past year’s public PDB deposits versus proprietary structures—and an overview of this curation workflow are presented in Figure 2. Our benchmarking dataset contains 113 antibody-antigen complexes (38 public, 75 proprietary) and 101 NANOBODY^®^ VHH-antigen complexes (28 public, 73 proprietary). We release all public PDB entries included in our benchmark to support evaluation of additional prediction methods.

### 4.3 DockQ Evaluation for Multi-Chain Epitopes

Accurate evaluation of antibody–antigen predictions requires careful handling of multi-chain epitopes, where the binding interface spans multiple antigen subunits. Applying DockQ (Mirabello and Wallner, 2024) naively introduces several failure modes: GlobalDockQ averages over irrelevant chain interfaces (antigen–antigen, heavy–light), diluting the binding signal and biasing scores even for correct predictions; per-chain evaluation fragments composite epitopes, so a prediction correctly docking the antibody at a subunit junction may score poorly against each chain individually; and homo-oligomeric antigens are sensitive to arbitrary chain labeling. DockQ v2 provides a `–mapping` flag that partially addresses chain-selection and label-ambiguity, but requires the correct mapping to be specified *a priori* and provides no mechanism to merge multiple antigen chains into a composite epitope entity, making it insufficient for automated benchmarking over a heterogeneous database where epitope composition is not known in advance.

We developed a custom evaluation methodology (Algorithm S3, Supplementary Section S4) that addresses each of these issues. Epitopes are classified as single- or multi-chain based on the fraction of interface contacts contributed by each antigen chain. For single-chain epitopes, DockQ is computed for each antigen chain independently and the maximum is reported; for multi-chain epitopes, all contributing chains are merged into a single composite entity before DockQ is computed, ensuring the metric evaluates the complete interface geometry. To handle labeling ambiguity, particularly for homo-oligomeric antigens, all sequence-compatible chain permutations are enumerated and the maximum DockQ across valid mappings is reported. For antibodies, VH and VL chains are merged into a single paratope entity prior to evaluation; for NANOBODY^®^ VHHs, the VHH chain is used directly.

## Abbreviations

Ab: antibody
Ag: antigen
ASU: asymmetric unit
C_*α*_: alpha carbon
CDR: complementarity-determining region
CDR-H3: heavy chain 3rd CDR
DockQ: docking quality score
Fab: fragment antigen-binding
FR: framework region
Fv: variable fragment
ipTM: interface predicted template modeling score
ML: machine learning
MSA: multiple sequence alignment
Nb: N-ANOBODY^®^ VHH
npy: NumPy array format
OF3p2: OpenFold-3 Preview2
PDB: Protein Data Bank
pLDDT: predicted local distance difference test
RF3: RosettaFold-3
SAbDab: Structural Antibody Database
SNAC-DB: Structural NANOBODY^®^ VHH and Antibody Complex Database
SOTA: state-of-the-art
TCR: T cell receptor
TM: template modeling
VH: variable heavy chain
VHH: variable heavy chain of heavy-chain-only antibody
VL: variable light chain.

## Supplementary Material

This article is accompanied by Supplementary Information containing detailed documentation of the SNAC-DB curation pipeline, benchmarking dataset construction, and analytical methods. The Supplementary Information includes:

- **Section S1:** Detailed SNAC-DB pipeline methodology, including cleaning and annotation procedures, immunoglobulin chain identification, VH-VL pairing logic, antigen assignment criteria, and implementation instructions.
- **Section S2:** Extended comparison with SAbDab, including curation capability analysis, structural diversity metrics, and a systematic interface composition analysis comparing epitope secondary structure, interface size, CDR contact distributions, and antigen sequence novelty between SNAC-DB and SAbDab.
- **Section S3:** Benchmarking dataset curation methodology, describing the four-stage filtering workflow (novel targets, novel epitopes, novel conformations, redundancy removal) with complete algorithmic specifications.
- **Section S4:** DockQ calculation methodology for multi-chain epitopes, including the CDR gate and contiguity filter, contact-based epitope classification, chain permutation handling, and differential treatment of single-vs. multi-chain cases.
- **Supplementary Figures S1–S8:** Pipeline schematics (S1–S4), interface diversity comparison plots (S5), antigen sequence novelty analysis (S6), benchmark structural novelty relative to the SNAC-DB training set (S7), and the impact of sample size on Boltz-2 performance using ipTM-based ranking versus oracle selection (S8).
- **Supplementary Algorithms S1–S3:** Formal algorithmic descriptions of the SNAC-DB curation pipeline (Algorithm S1), benchmarking dataset construction (Algorithm S2), and DockQ calculation for multi-chain epitopes (Algorithm S3).

Filename: `SNAC_DB_Supplementary_Information.pdf`

## Author Contributions

**Conceptualization:** A.G., B.M.R., Y.F.N., Y.Q., J.R.T., N.F.

**Methodology:** A.G., B.M.R., Y.Q., J.R.T.

**Software:** A.G., B.M.R., R.L.

**Validation:** A.G., B.M.R., R.L., J.R.T.

**Formal Analysis:** A.G., B.M.R., R.L., J.R.T.

**Investigation:** A.G., B.M.R., R.L., J.R.T.

**Resources:** N.F., Y.Q., Y.F.N., M.W.

**Data Curation:** A.G., B.M.R., R.L.

**Writing – Original Draft:** A.G., B.M.R.

**Writing – Review & Editing:** A.G., B.M.R., R.L., J.R.T., Y.F.N., M.W., Y.Q., N.F.

**Visualization:** A.G., B.M.R.

**Supervision:** A.G., N.F., Y.F.N., M.W., Y.Q.

**Project Administration:** A.G.

## Data Availability Statement

All code for the SNAC-DB curation pipeline is available under an open-source license at GitHub: https://github.com/Sanofi-Public/SNAC-DB. The ML-ready subset of public antibody, NAN-OBODY^®^ VHH, and TCR–antigen complexes—processed, trimmed, and formatted for immediate use in downstream model training and evaluation—can be downloaded from Zenodo: https://doi.org/10.5281/zenodo.15870002. Together, these resources provide a turnkey solution for both reproducing our curation workflow and leveraging the enriched structural database in new machinelearning and bioinformatics applications.

## Acknowledgments

We are grateful to Madhumati Sevvana, Ryan Gavin Casner, Prof. Thi Vo (Johns Hopkins University) for insightful discussions, and to the Sanofi Structural Biology teams for access to proprietary structures used in our benchmarking. We thank the BioAIM team—particularly Sabyasachi Dasgupta, Hervé Minoux, Andrew Phillips, Fred Fu, and Toni Anev—for providing the compute infrastructure that made this work possible. We also appreciate the encouragement and guidance of our leadership, Rebecca Sendak and Thorsten Schmidt.

## Supplementary Information

This supplementary information provides a detailed breakdown of all steps involved in constructing the Structural NANOBODY^®^ VHH and Antibody (VH-VL) Complex Database (SNAC-DB) and the analysis components used throughout the study.

## S1 Methodology of SNAC-DB

This section outlines the design and implementation of the SNAC-DB curation pipeline. The pipeline consists of three primary stages, each responsible for a distinct component of the curation process. We describe each stage in detail, including underlying assumptions and the rationale behind key design choices. For instructions on executing the pipeline, see the implementation section below.

Figure S1 illustrates the SNAC-DB pipeline applied to PDB structure 1FE8, providing a high-level overview of the processing at each stage.

**Figure S1:**
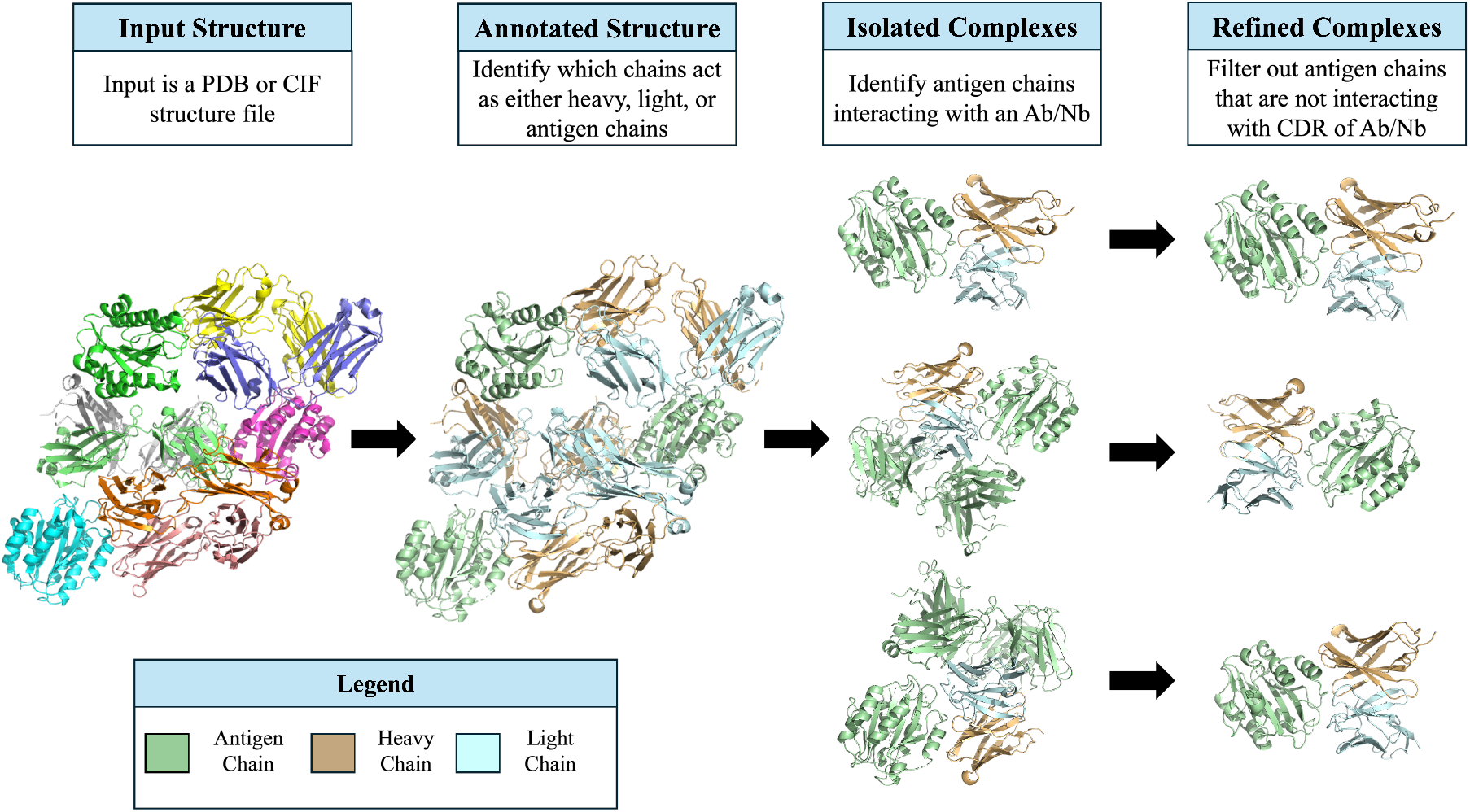
SNAC-DB pipeline applied to crystal structure 1FE8. The far-left panel displays the original input structure. In Step 1, the pipeline identifies heavy, light, and antigen chains (color-coded in the legend). In Step 2, antigens interacting with each ligand (Ab or Nb) are identified and separated into distinct complexes (third column). In the final step, the pipeline filters out antigen chains that do not directly interact with the ligand’s CDR regions, yielding biologically relevant complexes.

### S1.1 Cleaning and Annotating Structure Files

The first stage of the SNAC-DB pipeline prepares raw structural data for downstream processing, including resolving missing residues, standardizing nomenclature, and annotating each chain as heavy (VH), light (VL), or antigen (Ag). These annotations are assigned using a combination of open-source tools—including a modified version of ANARCI—and custom logic built into the pipeline.

The outputs of this stage include cleaned PDB structure files, metadata stored in NPY format, and a summary CSV file containing information required for complex identification in the next stage. A high-level schematic of this stage is shown in Figure S2.

**Figure S2:**
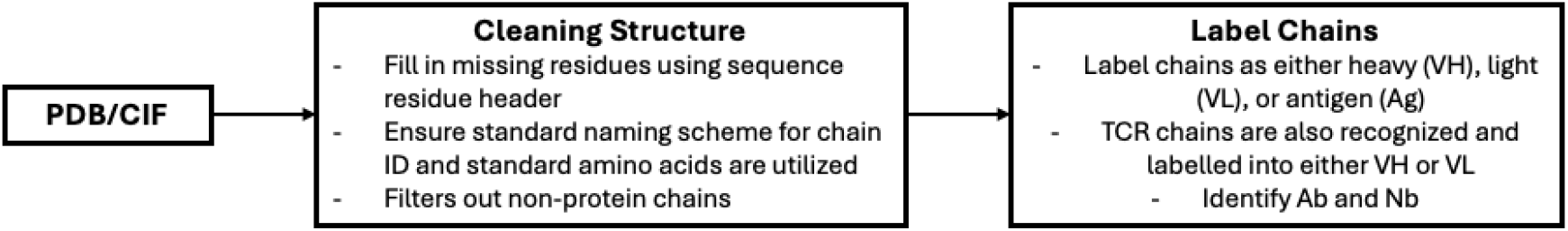
Overview of the cleaning and annotation stage in the SNAC-DB pipeline. Each input structure file is first standardized: chain identifiers follow standard naming conventions, non-protein chains are filtered out, and non-standard amino acids are excluded. Missing residues are resolved where possible using sequence information from the structure file header. ANARCI is then used to classify chains as heavy (VH), light (VL), or antigen (Ag). Antibodies (VH-VL pairs) and NANO-BODY^®^ VHHs (single VH chains) are identified based on contact interactions between the VH and VL chains.

#### S1.1.1 Residue and Naming Cleanup

Due to experimental limitations, some PDB structures contain unresolved residues—regions where residue identities and/or atomic coordinates are missing. These are typically represented as sequence gaps or ambiguous residues in the coordinate section; however, the full sequence is often available in the structure file header.

SNAC-DB uses this header sequence data to reconstruct missing residues where possible, improving structural completeness. The pipeline also enforces consistent naming conventions for residues and chain identifiers while preserving original chain IDs in output files for traceability and alignment with external references.

#### S1.1.2 Immunoglobulin Chain Annotation Using ANARCI

To assign immunoglobulin chain types, the pipeline uses ANARCI, which performs multiple sequence alignment (MSA) against an internal database. We employ a modified version of ANARCI that handles sequences containing unresolved residues. Chain identity is determined based on the first MSA match, classifying chains as either heavy (VH) or light (VL). ANARCI enables isolation of the variable region (Fv) from heavy and light chains, which are then stored as potential ligands. Untrimmed versions of VH and VL chains, along with all other chains not identified as immunoglobulin variable domains, are labeled as antigen (Ag) chains.

Variable regions are isolated for ligand representation because they drive antigen binding and are most relevant for structural modeling. The constant domain, which does not participate in binding, is excluded. Full chains are retained as potential antigens because the pipeline allows immunoglobulin chains from one complex to serve as antigens in another complex.

#### S1.1.3 Distinguishing Antibodies and NANOBODY^®^ VHHs

To differentiate between antibodies (paired VH and VL chains) and NANOBODY^®^ VHHs (unpaired VH chains), SNAC-DB uses contact-based heuristics derived from structural proximity. This workflow is illustrated in Figure S3.

The pipeline examines specific structural regions that contribute to VH–VL pairing, including complementarity-determining region 3 (CDR3), which plays a central role in the interaction interface, and framework regions FR2 and FR3, which are directly involved in interchain contacts. Portions of FR1 and FR4 are also evaluated; although these regions do not directly maintain the complex, they help reduce false positives arising from crystal packing artifacts or atypical VH–VL orientations. Despite conformational diversity among VH–VL complexes and the challenge of resolving replicate interactions in crystal structures, consistent spatial proximity across these regions enables robust and sensitive classification.

To evaluate contacts, the algorithm computes C_*α*_–C_*α*_ distance matrices between heavy and light chains using only residues within the selected FR and CDR regions. Three distance cutoffs—8 Å, 10 Å, and 12 Å—are applied sequentially to ensure flexible yet accurate detection of valid VH–VL interactions.

If valid contacts are identified, the VH-VL pair is classified as an antibody. Otherwise, the VH chain is treated as a NANOBODY^®^ VHH.

**Figure S3:**
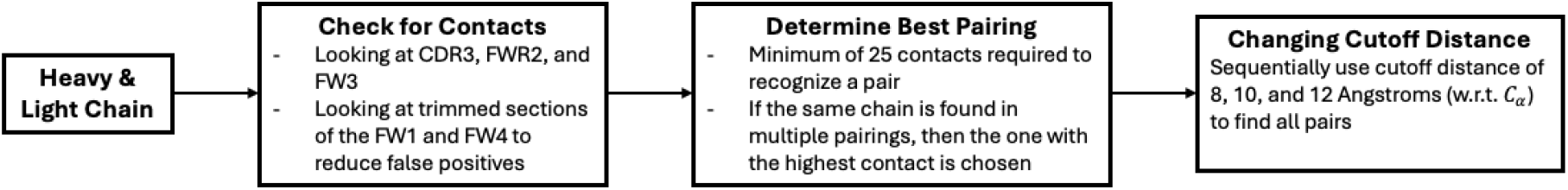
Workflow for identifying VH-VL pairs. Pairing is determined using CDR3, FWR2, and FWR3, which are directly responsible for VH-VL binding interactions. Sections of FWR1 and FWR4 are also evaluated to reduce false positives while maintaining sensitivity. These criteria are applied at cutoff distances of 8, 10, and 12 Å to ensure comprehensive detection of all VH-VL pairs.

#### S1.1.4 Outputs

Upon completion of cleaning and annotation, SNAC-DB produces:

- Cleaned PDB files with standardized chain labels and resolved residues
- NumPy (.npy) files containing dictionaries with keys VH, VL, and Ag, where each key maps to a sub-dictionary of chain IDs containing:
  - Amino acid sequence
  - Atom37 coordinates
  - B-factors
  - IMGT numbering (for VH and VL only)
- Summary CSV files with metadata for each structure:
  - Experimental method and resolution
  - Deposition and release dates
  - Chain IDs for VH, VL, and Ag chains

### S1.2 Identifying Complexes

This stage detects potential antigens interacting with identified ligands from the previous step. A permissive interaction definition is applied to minimize false negatives and ensure biologically relevant complexes are not missed. Each ligand—either an antibody (VH-VL) or NANOBODY^®^ VHH—is evaluated for proximity-based CDR contacts with potential antigen chains.

#### S1.2.1 Interaction Criteria

To maximize sensitivity during this initial filtering stage, the following criteria identify potential antigen chains:

- At least one antigen residue must lie within 12 Å (C_*α*_–C_*α*_ distance) of the ligand’s CDRs
- The contact may occur in any CDR of the heavy or light chain

Although the 12 Å threshold is relatively generous, it accounts for resolution variability, minor structural deviations, and potential crystal packing distortions. This relaxed criterion is justified by two factors: (1) stricter filtering is applied in the subsequent refinement stage, and (2) the copresence of ligands and antigens in experimental structures typically implies meaningful interaction, even if loosely resolved.

#### S1.2.2 Complex Renaming and Standardization

Once ligand and interacting antigen chains are identified, the pipeline constructs a new complex and applies standardized chain labeling:

- Heavy chains (VH) are relabeled as H
- Light chains (VL), when present, are relabeled as L
- Antigen chains are assigned IDs from A to Z (skipping H and L); if more than 24 antigens are present, IDs from 0–9 and a–z are used

To maintain traceability, original chain IDs are recorded in the REMARK section of each PDB file and included in the structure title, enabling users to map curated complexes back to source chains.

#### S1.2.3 Outputs

Each identified complex is saved in two formats:

- **PDB file**: Cleaned, relabeled complex structure
- **NumPy (.npy) file**: Dictionary with chain IDs as keys, where each maps to a sub-dictionary containing:
  - Chain type (VH, VL, or Ag)
  - IMGT numbering (for VH or VL only)
  - Amino acid sequence
  - Atom37 coordinates
  - B-factors
  - Original chain ID
  - Pair key (when applicable), indicating the chain ID of a paired VH or VL acting as antigen

Additionally, a summary CSV file is generated containing structural metadata from the original structure (PDB ID, experimental method, resolution, classification, deposition and release dates) and updated annotations for each complex:

- New and original chain IDs for VH, VL (if present), and Ag
- Antigen pairing relationships, if identified
- Complex description in comment field

### S1.3 Refining Complexes

This stage applies stricter validation criteria to complexes identified in the previous step, verifying genuine binding interfaces and eliminating false positives from crystal packing or incidental proximity. A schematic of antigen-ligand interaction determination (from the **Identifying Complexes** and **Refining Complexes** stages) is shown in Figure S4.

**Figure S4:**
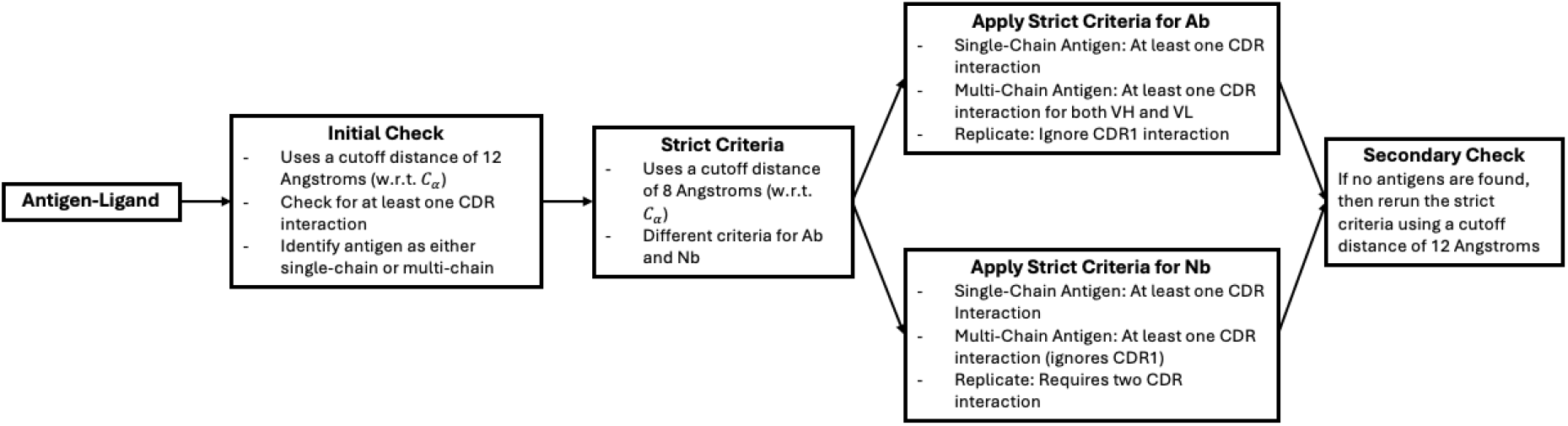
Workflow for determining antigen-ligand interactions. The pipeline employs lenient criteria in the initial identification stage (12 Å cutoff), followed by stringent validation in the refinement stage (8 Å cutoff) that depends on antigen type (single-chain or multi-chain) and ligand type (Ab or Nb). If no antigens pass 8 Å validation, a rescue pass applies the same validation rules at 12 Å.

#### S1.3.1 Contact Criteria

An 8 Å C_*α*_–C_*α*_ distance cutoff serves as the primary validation threshold. Specific criteria depend on ligand type (antibody or NANOBODY^®^ VHH) and whether the antigen is single-chain or multichain.

For multi-chain antigens, when one chain meets the 8 Å interaction criteria, other chains identified during the 12 Å identification stage are retained to preserve the complete functional epitope. In special cases, when only one chain of a VH-VL pair contacts the antigen, the non-contacting chain is explicitly included, reflecting their cooperative structural binding properties.

When the antigen shares the same VH or VHH sequence as the ligand (potential crystal symmetry artifacts or weak cognate pairs), additional stringent criteria are applied. For antibody complexes, contacts must occur in both CDR2 and CDR3 of both VH and VL chains. For NANOBODY^®^ VHH complexes, at least two CDR contacts must be present. This filters spurious inter-ligand crystal contacts while preserving legitimate cases where antibodies or NANOBODY^®^ VHHs act as antigens.

#### S1.3.2 Antibody Complexes

- **Single-chain antigens:** Must have at least one CDR contact with either the heavy or light chain at 8 Å
- **Multi-chain antigens:** Both VH and VL chains must have CDR contacts with the antigen complex at 8 Å. When one chain of the multi-chain antigen meets the interaction criteria, other chains identified at 12 Å are retained to preserve epitope integrity (with exceptions when a VH-VL pair acts as part of the antigen)

The dual-contact requirement for multi-chain antigens verifies proper antibody orientation toward the antigen, serving as a safeguard against false-positive interactions from partial or indirect proximity in multimeric structures.

#### S1.3.3 NANOBODY^®^ VHH Complexes

- **Single-chain antigens:** Must have at least one CDR contact with the NANOBODY^®^ VHH at 8 Å
- **Multi-chain antigens:** Must have at least one CDR contact (excluding CDR1) with the NANOBODY^®^ VHH at 8 Å. When one chain meets the criteria, other chains identified at 12 Å are retained to preserve the complete epitope

The stricter criteria for multi-chain antigens (excluding CDR1) reflect the expectation of a more extensive binding interface, ensuring retained interactions are biologically meaningful.

#### S1.3.4 Rescue Pass

If no valid antigen chains are identified after applying 8 Å validation criteria, and the PDB entry contains additional chains beyond VH–VL or VHH, the pipeline performs a rescue pass applying the same validation rules with a 12 Å C_*α*_–C_*α*_ cutoff instead of 8 Å.

This fallback recovers complexes with poorly resolved interactions, low-resolution structures, or atypical binding geometries that may otherwise be filtered out. The co-presence of ligand and antigen chains in experimental structures strongly suggests intentional complex formation, even if the interface is partially obscured or distorted by crystallization conditions.

#### S1.3.5 Outputs

Outputs for this stage match those from the previous step, with updated complex names reflecting filtered antigen chains. Structure and metadata files are generated with the key differences being: (1) some antigen chains have been removed, resulting in new structure filenames, and (2) metadata reflects the updated set of interacting antigen chains.

#### S1.3.6 Redundancy Removal

To eliminate near-identical structures arising from multiple biological assemblies or asymmetric units within the same PDB entry, we employed FoldSeek multimersearch to identify redundant structures (TM-score > 0.999). When multiple structures from the same PDB ID were flagged as redundant, we retained only the highest-priority structure based on a weighted scoring system: priority = (total residues × 500) - (missing residues × 100) + assembly bonus - assembly number, where biological assemblies received a +1000 bonus and asymmetric units received +500. This approach ensures retention of the most complete, highest-quality structure while eliminating redundant entries that provide no additional structural information. For example, if PDB entry 7ABC contained three structures with TM-scores > 0.999, we retained only the structure with the most residues, fewest missing C_*α*_ atoms, and preference for biological assembly over asymmetric unit.

### S1.4 Implementing the SNAC-DB Pipeline

The SNAC-DB pipeline consists of four sequential components:

1. **Processing and Annotating Structure Files:** Cleans the input structure files and assigns chain labels (VH, VL, Ag).
2. **Identifying Complexes:** Detects potential immunoglobulin complexes based on CDR interactions.
3. **Refining Complexes:** Applies stricter interaction criteria to filter out non-interacting antigen chains.
4. **Redundancy Removal:** Eliminates near-identical structures arising from multiple biological assemblies or asymmetric units within the same PDB entry.

Each component generates a directory containing curated structure files and a corresponding summary CSV file. The final curated dataset, produced by the refinement step, is typically of most interest. These files can be found in the directory named **input_directory_curated** or **in-put_directory_filter**, depending on the optional parameters inputted when running the pipeline. The complete pipeline logic, including all filtering stages and output generation, is formalized in Algorithm S1.

Note: Running the pipeline multiple times in the same location will overwrite existing output files. To preserve previous results, users should rename the output directories or execute the pipeline in a new directory with a copied version of the input files.

In this section we only provide a brief explanation of how to utilize the pipeline; however, a more detailed explanation can be found on the SNAC-DB GitHub page.

#### S1.4.1 Implementing the Pipeline Using the Bash Script

The recommended approach is to use the provided bash script **data_curation.sh**, which automatically executes all three pipeline stages and requires only the path to the directory containing input structure files.

Two optional parameters are available: (1) whether to create intermediate files (structure and metadata files from each pipeline stage), and (2) whether to eliminate redundant complexes from the final curated dataset. By default, intermediate files are not created and redundant complexes are eliminated.

#### S1.4.2 Output File Naming Convention

Each curated complex is saved as .pdb and .npy files using a consistent naming scheme. An example filename is **8FSL-ASU1-VHH_B-Ag_C**, composed of the following elements:

- **PDB ID:** Original structure identifier (e.g., 8FSL)
- **Assembly tag:** Bioassembly number (e.g., ASU1); ASU0 denotes the asymmetric unit when no assembly is specified
- **Complex descriptor:** Ligand type (VHH for NANOBODY^®^ VHH, VH-VL for antibody) followed by original chain IDs for ligand and antigen

This format ensures unique identification and traceability to source structures.

#### S1.4.3 Handling Replicated Chains

When a chain contains multiple sub-chains (e.g., a chain containing two NANOBODY^®^ VHHs bound to the same antigen), the default naming scheme may produce identical filenames. To prevent overwriting, the pipeline appends a replicate identifier, such as **6UL6-ASU1-VHH_B-Ag_A-replicate0** and **6UL6-ASU1-VHH_B-Ag_A-replicate1**.

The replicate integer corresponds to the sub-chain partition and may vary between pipeline runs, as sub-chain assignment is not strictly deterministic. However, the remainder of the filename remains consistent, preserving contextual information about the complex and its origin.

## S2 Comparison Between SNAC-DB and SAbDab

This section compares the Structural NANOBODY^®^ VHH and Antibody (VH-VL) Complex Database (SNAC-DB) to the Structural Antibody Database (SAbDab) across two dimensions: curation capability and structural diversity.

### S2.1 Curation Capability

To ensure fair comparison, both pipelines were evaluated on the same input structures. As of March 31, 2025, SAbDab had curated 18,717 entries (both complexes and ligand-only structures) from 9,506 PDB structures. However, SNAC-DB extracts only protein–protein complexes involving heavy chains (antibodies or NANOBODY^®^ VHHs) and excludes non-protein antigens and unbound light chains. For parity, we applied these same filters to SAbDab’s dataset, removing entries lacking antigen chains or representing isolated VL chains. After filtering, 16,964 complexes remained across 8,914 PDB structures.

We then excluded 437 complexes that SNAC-DB could not process properly. Failures occurred due to download errors, ANARCI misidentifying ligand chains, or structure file parsing issues. Notably, some PDB entries had bioassembly files from which SNAC-DB successfully extracted valid complexes; however, since SAbDab processes only asymmetric units, we limited comparison to structures both pipelines could process. This yielded a final matched dataset of 16,527 complexes from 8,670 PDB entries.

### S2.2 Structural Diversity

To assess structural diversity, we expanded the analysis beyond the matched dataset.

For SNAC-DB, we curated all PDB entries deposited on or before March 31, 2025, retaining only those with at least one valid complex, then eliminated redundant complexes (defined as structurally identical). This yielded 15,523 curated complexes. For SAbDab, we used the filtered dataset described above, excluding structures lacking protein antigen chains, resulting in 13,919 curated complexes.

To ensure consistency, the Fc region (constant domain) was trimmed from each ligand in both datasets prior to comparison. We then clustered each dataset independently using Foldseek-Multimer at TM-score thresholds of 0.60, 0.70, 0.80, 0.90, and 0.95. The resulting cluster counts were:

- **SNAC-DB:** 2,580; 3,432; 4,250; 5,348; 6,659
- **SAbDab:** 1,873; 2,502; 3,179; 4,001; 5,047

This comparison quantifies the structural redundancy and diversity of complexes identified by each pipeline.

### S2.3 Interface Composition Analysis

To characterize what SNAC-DB adds beyond SAbDab at the level of physical interfaces rather than cluster counts, we conducted a systematic comparison of interface composition across both databases. Contacts are defined as any heavy atom within 5 Å of any heavy atom on the opposing side, applied identically to both datasets. This stringent any-atom cutoff ensures that observed differences in interface composition reflect genuine structural variation between the databases rather than artefacts of contact definition: a loose cutoff could artificially inflate the number of contacted residues or chains, but a 5 Å any-atom threshold requires physical proximity and cannot be gamed by threshold choice. Secondary structure was assigned with mkdssp (Hekkelman et al., 2025) and collapsed into three classes: helix (H/G/I), strand (E/B), and loop (T/S/space). mkdssp succeeded on 99.3% of SAbDab structures and 99.5% of SNAC-DB structures; remaining structures are excluded only from secondary-structure analyses.

#### Multi-chain epitope prevalence

The most striking difference between the two databases is in antigen chain multiplicity (Figure S5, top left). SNAC-DB contains 33.5% multi-chain antigen complexes versus 15.3% in SAbDab, more than double the rate, reflecting SNAC-DB’s explicit enumeration of all antigen chains within contact distance of the paratope. Multi-chain epitopes represent qualitatively distinct binding problems: the antibody/NANOBODY^®^ VHH must recognize a quaternary structure surface assembled from two or more protein subunits, and a prediction model must correctly position the binder relative to both chains simultaneously.

**Figure S5:**
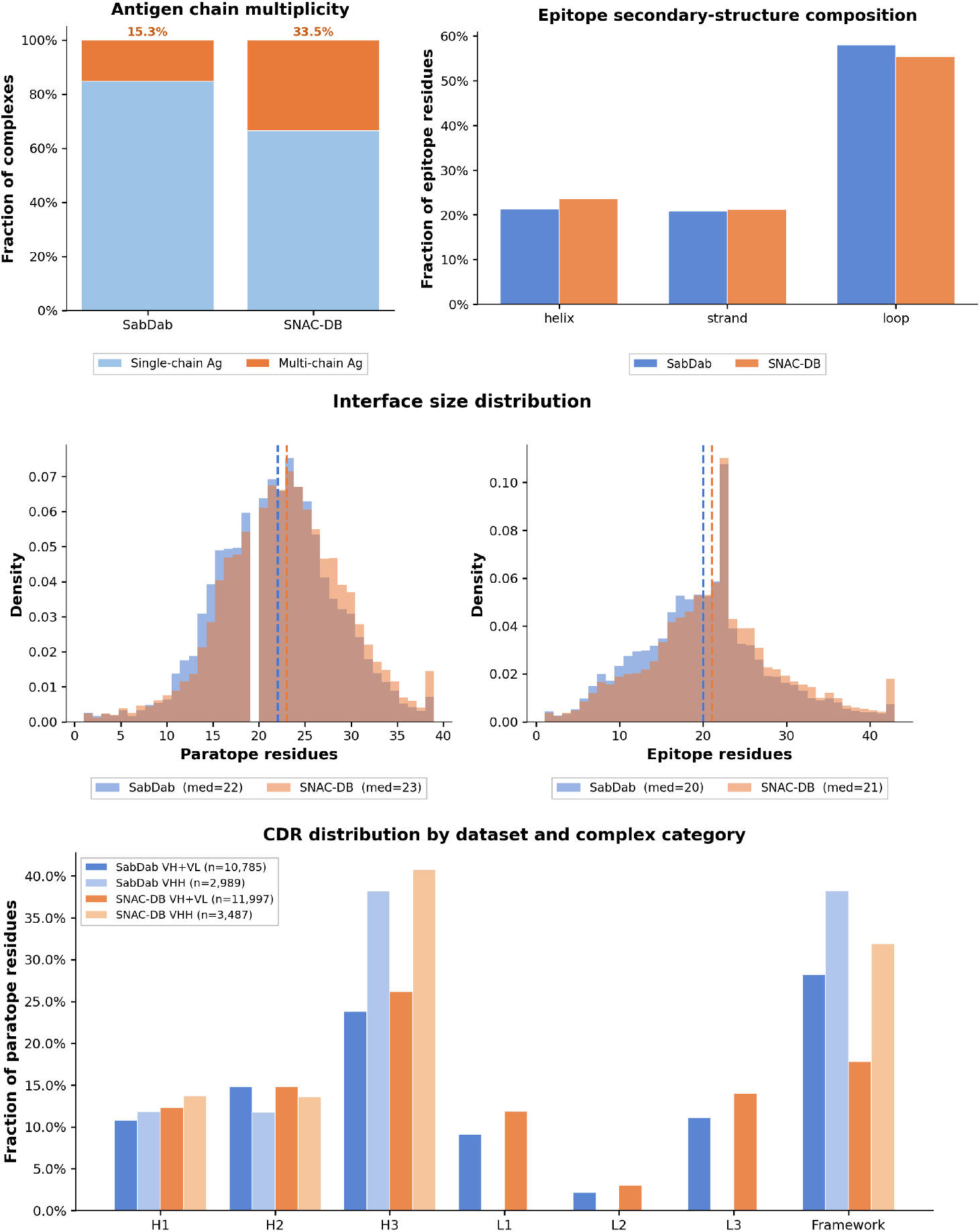
Interface composition comparison between SNAC-DB and SAbDab. **(Top left)** Fraction of complexes with single-vs. multi-chain antigens. SNAC-DB contains more than twice the proportion of multi-chain antigen complexes (33.5% vs. 15.3%), reflecting its biological-assembly-based curation. **(Top right)** Secondary-structure composition of epitope residues (DSSP-assigned). SNAC-DB shows a more balanced distribution, with a modestly lower loop fraction (55.3% vs. 58%) and corresponding increases in helix and strand contacts. **(Middle)** Interface size distributions (5 Å any-atom cutoff) for paratope (left) and epitope (right) residues. SNAC-DB shows modestly larger interfaces (paratope median: 23 vs. 22; epitope median: 21 vs. 20) with a heavier right tail, consistent with its larger multi-chain antigen fraction. **(Bottom)** Paratope CDR contact distribution broken down by antibody class (VH+VL and VHH) within each dataset. SNAC-DB shows a modestly more balanced distribution across CDR loops in both classes, with lower CDR-H3 dominance and higher framework region contacts, particularly for NANOBODY^®^ VHHs.

#### Epitope secondary-structure composition

SNAC-DB epitopes show a more balanced secondarystructure distribution than SAbDab: the loop fraction decreases from 58% to 55.3%, while helix and strand contacts increase correspondingly (Figure S5, top right). Although the differences are modest, they are consistent in direction across all three classes, indicating that SNAC-DB proportionally enriches non-loop epitope contacts relative to SAbDab. This is consistent with SNAC-DB’s inclusion of more multi-subunit antigens, which tend to present more regular secondary-structure elements at subunit interfaces. Models trained on SAbDab alone are therefore slightly over-exposed to loop-dominated epitopes relative to the broader distribution of antigen surface types captured by SNAC-DB.

#### Interface size

SNAC-DB interfaces are modestly larger than SAbDab (paratope median: 23 vs. 22 residues; epitope median: 21 vs. 20 residues; Figure S5, middle), with SNAC-DB showing a heavier right tail. Both differences are statistically significant at the large sample sizes involved (*p <* 10^−27^, Mann–Whitney U), though the practical magnitude is small. The right-tail enrichment is consistent with SNAC-DB’s larger multi-chain epitope fraction: composite epitopes spanning two antigen subunits are inherently larger than single-chain ones.

#### CDR contact distribution by antibody class

Across both antibody classes, SNAC-DB shows a higher fraction of paratope contacts contributed by CDR loops and a correspondingly lower framework fraction relative to SAbDab (Figure S5, bottom). For conventional antibodies (VH+VL), CDR-H3 contributes slightly more in SNAC-DB (∼26% vs. ∼23%) while the framework fraction is lower (∼17% vs. ∼27%). A similar pattern holds for NANOBODY^®^ VHHs, where SNAC-DB VHH complexes show higher CDR-H3 contribution (∼40% vs. ∼38%) and lower framework contacts (∼32% vs. ∼38%). Both databases consistently show a higher framework fraction for VHH complexes than for VH+VL, as expected given that NANOBODY^®^ VHHs rely more heavily on framework residues to compensate for the absence of light-chain CDRs. Within each antibody class, CDR loop distributions are otherwise broadly consistent between the two databases, confirming that SNAC-DB does not introduce systematic biochemical biases in paratope composition.

#### Antigen sequence novelty

To quantify how much of SNAC-DB’s antigen sequence space is absent from SAbDab, we used MMseqs2 `easy-search` to compute the maximum sequence identity of each SNAC-DB antigen chain against the full SAbDab antigen chain database (19,609 SNAC-DB chains searched against all SAbDab chains). A complex is counted as novel at threshold *T* if any of its antigen chains has maximum SAbDab identity strictly below *T* ; because the novel-chain set can only grow as *T* increases, this metric is monotonically non-decreasing by construction (Figure S6). At the stringent 30% identity threshold (essentially unrelated sequences), 2.5% of SNAC-DB complexes (*n*=358) carry an antigen chain with no detectable SAbDab homologue. At 90% identity, 9.6% of complexes (*n*=1,385) contain an antigen chain not well-represented in SAbDab, representing cases where training on SAbDab alone would provide no direct supervision signal for the antigen side of the interface.

##### Algorithm S1

SNAC-DB Data Curation Pipeline

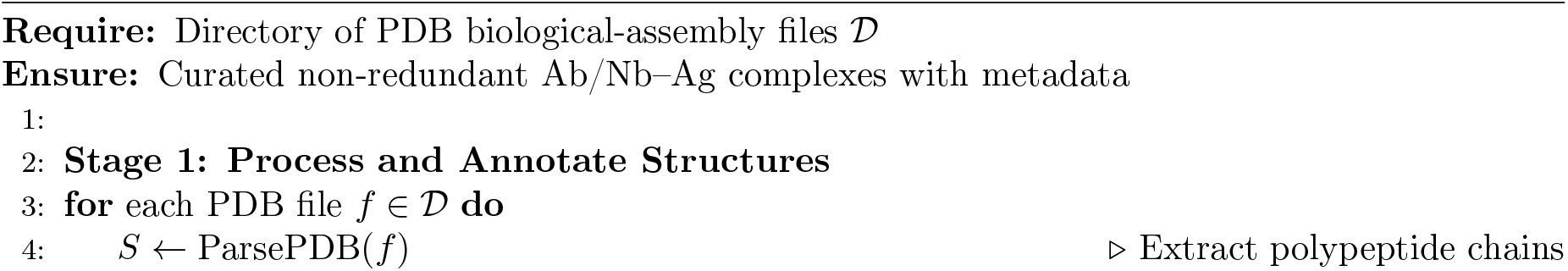
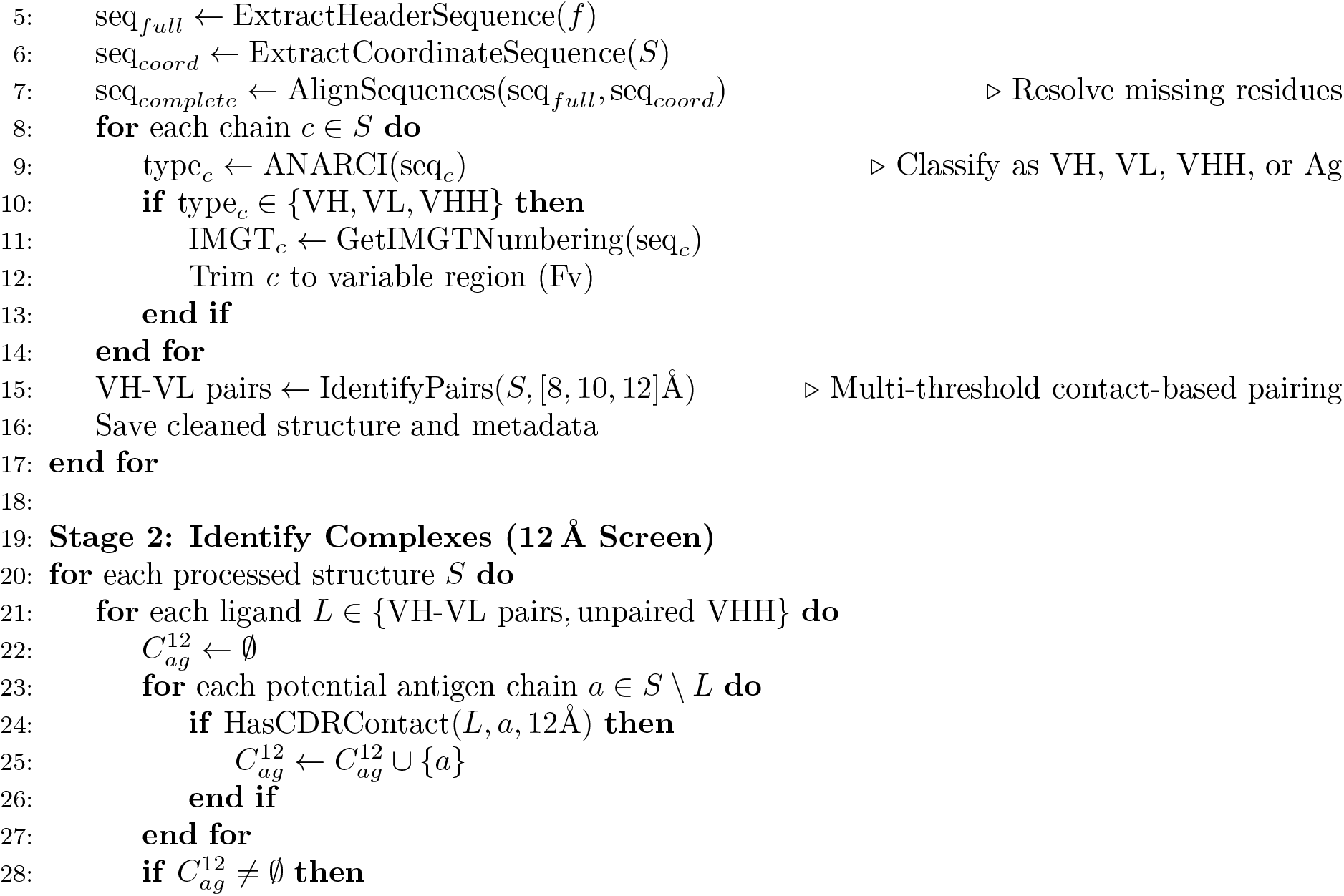
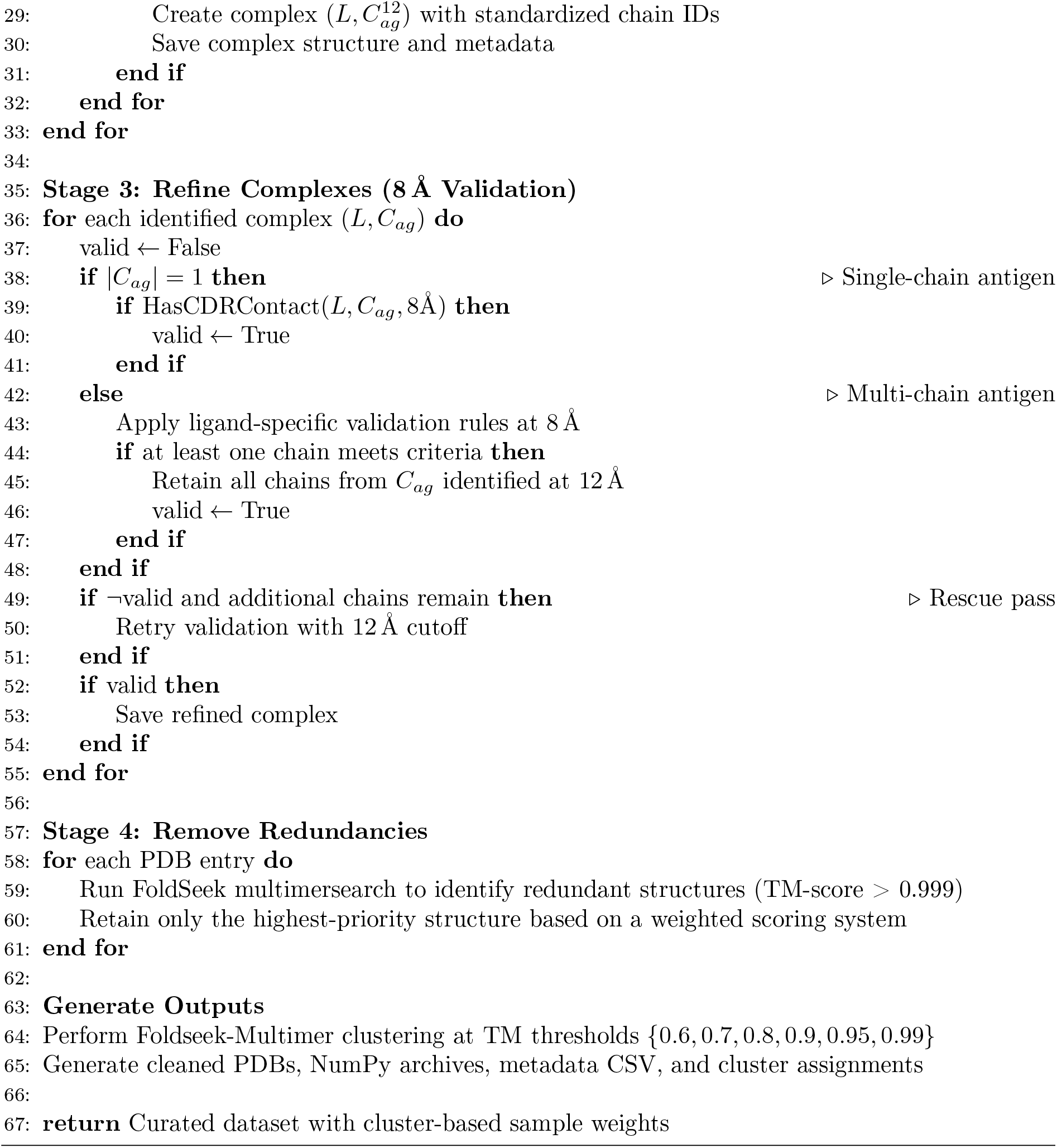

**Figure S6:**
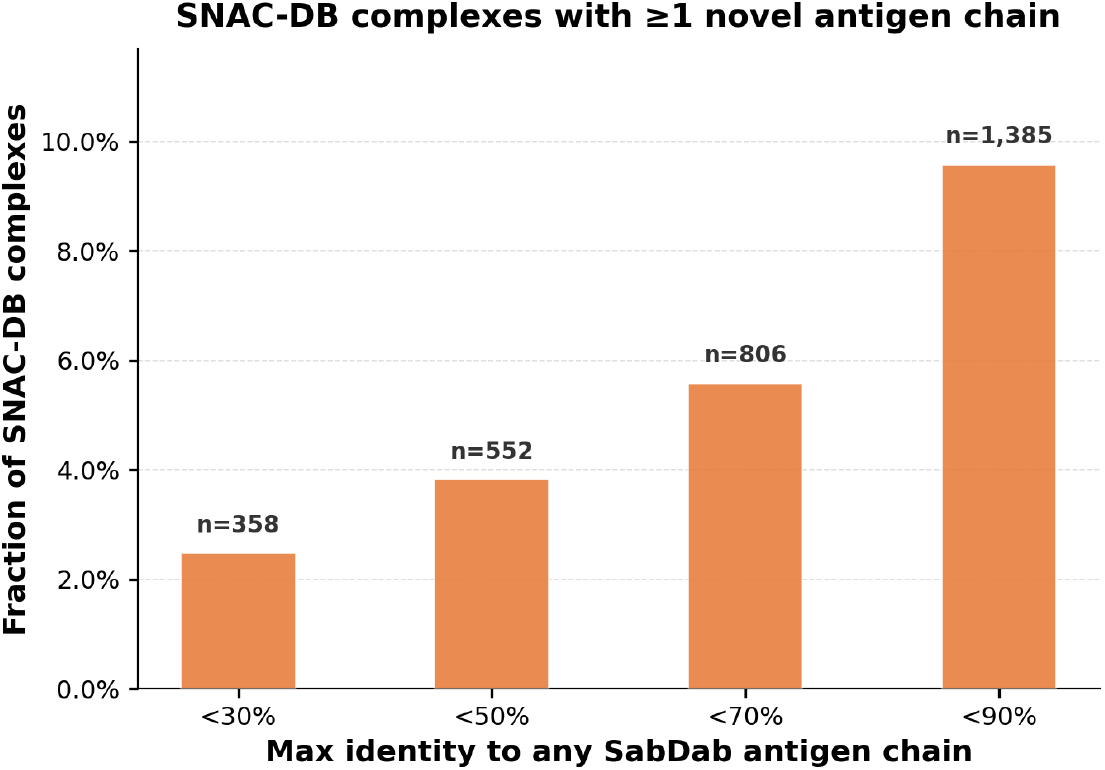
Antigen sequence novelty of SNAC-DB relative to SAbDab (search-based, monotone). For each SNAC-DB antigen chain, the maximum sequence identity to any SAbDab antigen chain was computed using MMseqs2 `easy-search` (19,609 SNAC-DB chains queried against all SAbDab chains). A complex is counted as novel at threshold *T* if any of its antigen chains has maximum SAbDab identity strictly below *T* . Because the novel-chain set can only grow as *T* increases, the resulting count is monotonically non-decreasing by construction.

## S3 Curation of Benchmarking Dataset

To evaluate model performance beyond documented training data, we curated a structurally diverse and non-redundant benchmark dataset composed of Ab/Nb complexes intentionally dissimilar to models’ publicly documented training sets. The goal is to challenge models with structures from regions of structural space designed to minimize overlap with known training data, thereby assessing real-world predictive performance.

The query dataset, used as the benchmark set, was assembled from approximately 600 curated proprietary structures and 400 curated immunoglobulin complexes from the PDB deposited after May 30, 2024. The reference dataset used for filtering contains a proprietary test set ( 100 curated complexes) and 14,000 curated immunoglobulin complexes from the PDB deposited before May 30, 2024 (SNAC-DB training set). The high-level overview is illustrated in Figure 2.

To ensure valid structural comparisons, we applied the following preprocessing steps:

- Retained only complexes with at least one polypeptide antigen chain ≥65 residues to avoid misleading similarity from small fragments
- Excluded complexes exhibiting weak cognate pair interactions—typically crystal packing artifacts between ligand replicates
- For PDB-derived structures, used only bioassembly models with resolution ≤4.0 Å to ensure structural reliability
- Performed manual inspection as final quality control, removing entries with misfolded chains or apparent modeling errors

Pairwise structural comparisons were performed using Foldseek (Kim et al., 2025), which supports both global single-chain and multi-chain structural alignment. Foldseek returns two Template Modeling (TM) scores per comparison, normalized by query and reference structures respectively. To mitigate size bias (e.g., from truncated or extended antigens), we retained the maximum of the two TM scores as the similarity metric. In Stages 1 and 2, we employed single-chain comparisons by merging multiple chains into a single chain ID (‘Z’), enabling Foldseek’s easy-search mode to perform rapid global alignment and detect broad structural similarity at the antigen or whole-complex level. In Stages 3 and 4, we used Foldseek’s easy-multimersearch mode with original multi-chain structures to capture subtle conformational differences and relative chain orientations that single-chain comparisons would miss. The benchmark curation pipeline consists of four sequential stages, with structures failing each stage advancing to the next more stringent filter.

### S3.1 Stage 1: Novel Targets (Antigen-Level Filtering)

#### Goal

Identify complexes with structurally novel antigens.

##### Structure preparation

Each complex in both query and reference datasets is processed to extract only antigen chains (removing H and L chains). For complexes with multiple antigen chains, all chains are merged into a single chain with ID ‘Z’ to enable single-chain comparison.

##### Comparison method

Foldseek easy-search mode compares query antigens against reference antigens using sensitivity parameter -s 7.5, outputting query and target TM-scores.

##### Classification

For each query complex, we identify the highest TM-score (maximum of query-normalized and target-normalized scores) among all reference comparisons. Complexes with maximum TM-score *<* 0.6 are classified as novel targets and pass to the final benchmark. Complexes with TM-score ≥ 0.6 fail and proceed to Stage 2.

##### Rationale

This stage assumes that novel antigen scaffolds imply novel epitopes and binding modes. The 0.6 threshold ensures substantial structural divergence from known antigens.

### S3.2 Stage 2: Novel Epitopes (Whole-Complex Global Alignment)

#### Goal

Among complexes with known antigens, identify those with novel binding sites or ligand orientations.

##### Structure preparation

Failed complexes from Stage 1 are processed to merge all chains (H, L, and all antigen chains) into a single chain with ID ‘Z’, converting each complex into a single-chain representation for global structural alignment.

##### Comparison method

Foldseek easy-search compares these single-chain representations of query complexes against similarly prepared reference complexes.

##### Classification

Complexes with maximum whole-complex TM-score *<* 0.6 are classified as novel epitopes and pass to the final benchmark. Complexes with TM-score ≥ 0.6 fail and proceed to Stage 3.

##### Rationale

By comparing entire complexes as single chains, we detect differences in binding site location or antibody/NANOBODY^®^ VHH orientation relative to the antigen, even when antigen scaffolds are similar. The same 0.6 threshold maintains consistency with antigen-level filtering.

### S3.3 Stage 3: Novel Conformations (Multi-Chain Structural Filtering)

#### Goal

Detect subtle conformational differences in complexes with similar antigens and epitopes.

##### Structure preparation

Failed complexes from Stage 2 retain their original multi-chain structures without modification.

##### Comparison method

Foldseek easy-multimersearch aligns multiple chains simultaneously, preserving chain identities and relative orientations. The algorithm validates that:

- All chains of the shorter complex (fewer total chains) are aligned
- Heavy chains map to heavy chains (H → H)
- Light chains map to light chains (L → L)
- Comparisons lacking proper chain correspondence are excluded

##### Classification

Complexes with maximum TM-score *<* 0.85 are classified as novel conformations and pass to the final benchmark. Complexes with TM-score ≥ 0.85 fail and are considered highly similar to reference structures.

##### Rationale

The higher threshold (0.85 vs 0.6) reflects our goal of retaining complexes with subtle but meaningful conformational variations. Multi-chain alignment captures dynamic variations in CDR loop conformations, relative VH-VL orientations, and interface geometry that single-chain comparisons miss. Chain mapping validation ensures biologically meaningful alignments.

### S3.4 Stage 4: Redundancy Removal (Internal Benchmark Filtering)

#### Goal

Ensure the final benchmark contains structurally diverse, non-redundant complexes.

##### Structure preparation

All complexes that passed Stages 1–3 are pooled and compared against each other using their original multi-chain structures.

##### Comparison method

Foldseek easy-multimersearch performs self-comparison of the benchmark set, applying the same chain mapping validation as Stage 3. To prevent double-counting, once a complex is identified as similar to another (TM-score ≥ 0.85), it is added to an exclusion list for subsequent comparisons.

##### Classification

For each complex, we identify its highest TM-score against all other benchmark complexes (excluding self-comparisons). Complexes with maximum TM-score *<* 0.85 pass as unique representatives. Complexes with TM-score ≥ 0.85 to another benchmark complex are excluded as redundant.

##### Rationale

This final stage ensures the benchmark itself is structurally diverse, preventing over-representation of similar binding modes. The 0.85 threshold balances retaining meaningful conformational variants while removing near-duplicates.

### S3.5 Final Output and Summary Statistics

All complexes passing Stage 4 are combined with those that passed earlier stages to form the final benchmark dataset. A summary CSV file tracks each complex’s classification (novel target, novel epitope, novel conformation), closest reference match, TM-score to that match, and pass/fail status at each stage. This transparent documentation enables users to subset the benchmark based on specific evaluation criteria or analyze performance trends across different complexity levels. See Algorithm S2 for the complete curation workflow. Figure S7 validates the curation from two complementary perspectives: the left panel confirms that benchmark complexes are structurally novel relative to the SNAC-DB training set, with TM-scores consistent with the intended curation thresholds; the right panel shows that public and proprietary subsets are structurally dissimilar to each other, confirming that they probe distinct regions of structural space rather than representing overlapping coverage.

**Figure S7:**
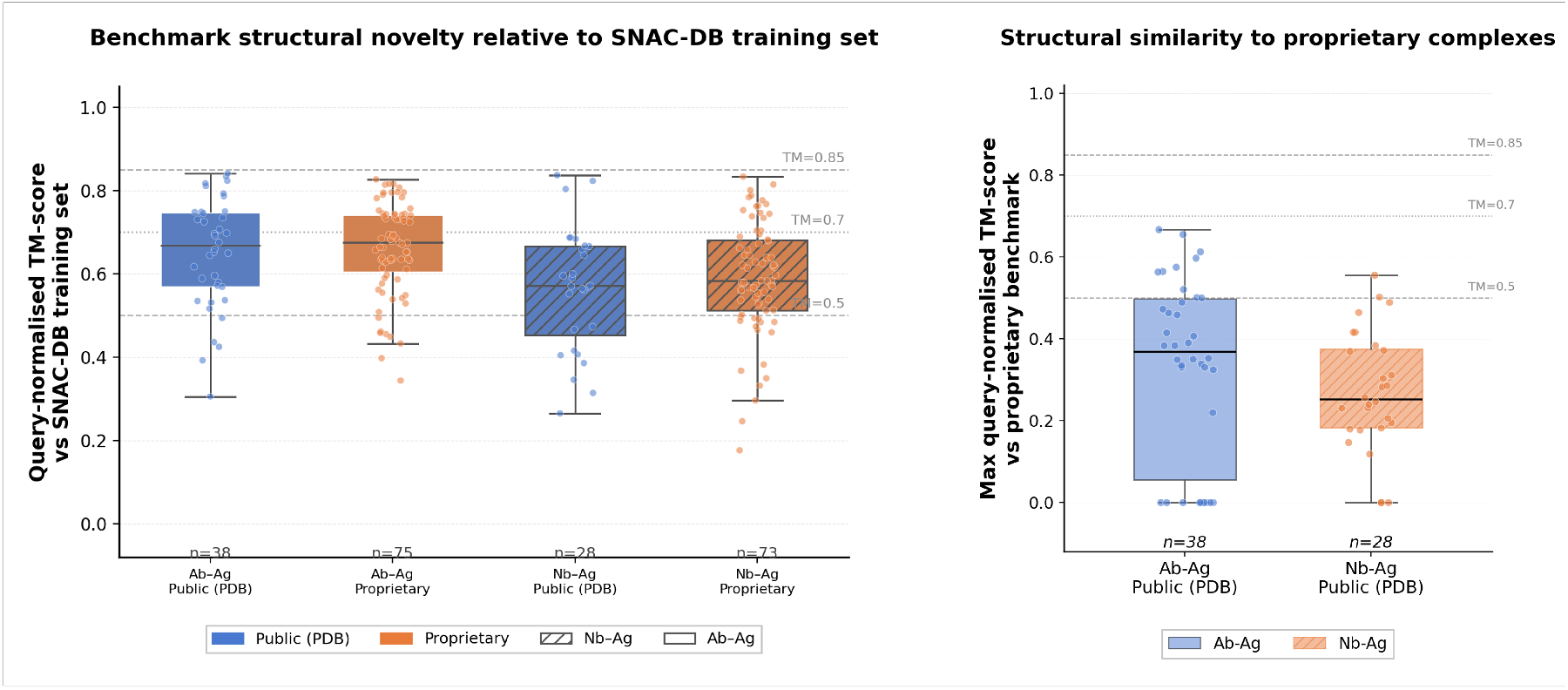
Structural novelty and diversity of the benchmark dataset. *Left:* Query-normalised TM-score of each benchmark complex against its nearest neighbour in the SNAC-DB training set (computed via FoldSeek Multimer-search), shown separately for Ab–Ag and Nb–Ag complexes and split by public (PDB) and proprietary origin. Lower TM-scores indicate greater structural novelty. Reference lines are drawn at TM=0.5, 0.7, and 0.85, corresponding approximately to the thresholds for different fold similarity and the novel conformation cutoff used in benchmark curation. *Right:* Maximum query-normalised TM-score of each public benchmark complex against any proprietary complex in the benchmark, for Ab–Ag and Nb–Ag separately. Low values indicate that public and proprietary complexes are structurally dissimilar to each other, confirming that the two subsets probe distinct regions of structural space.

#### Algorithm S2

Benchmarking Dataset Curation Pipeline

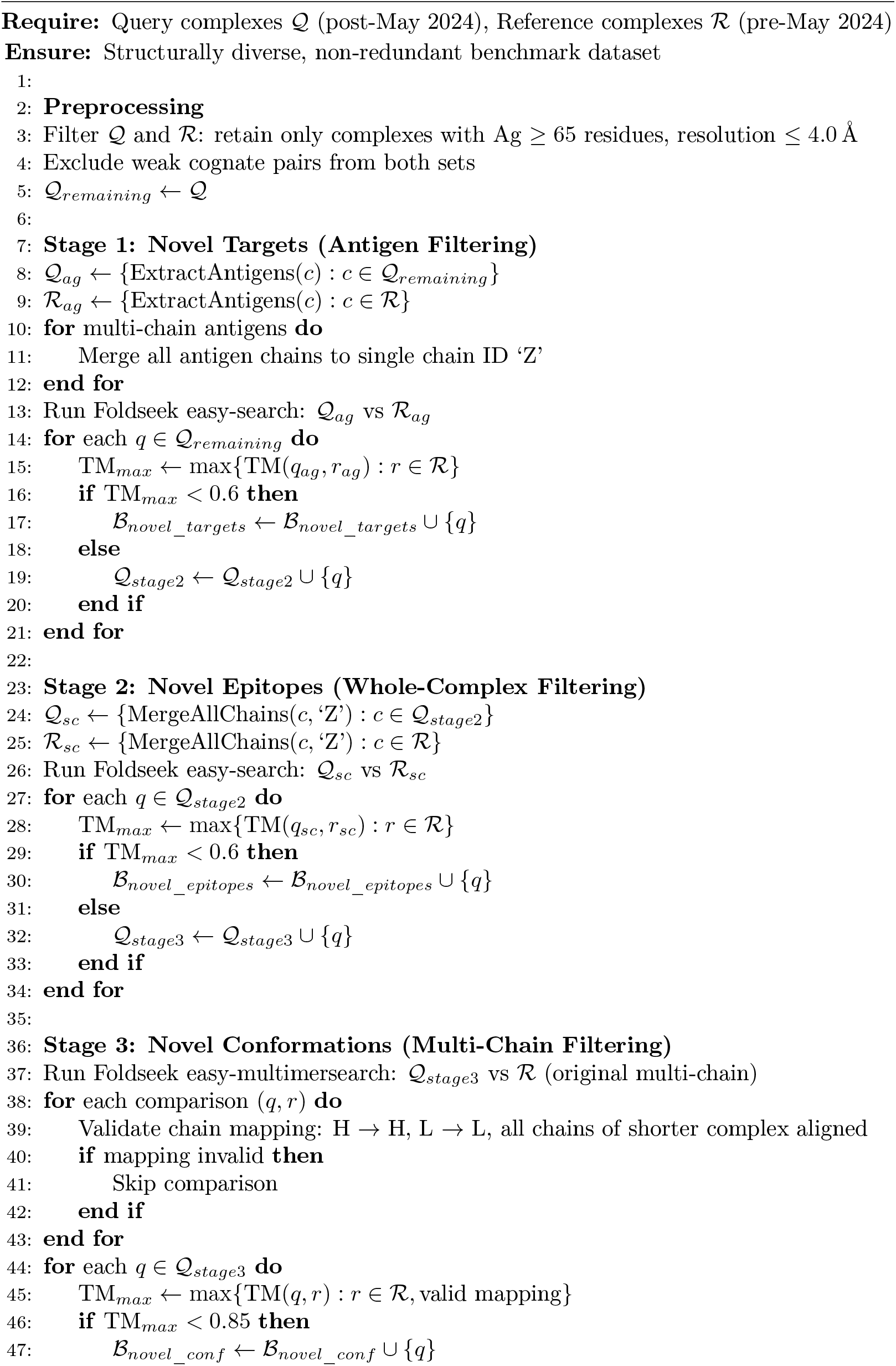
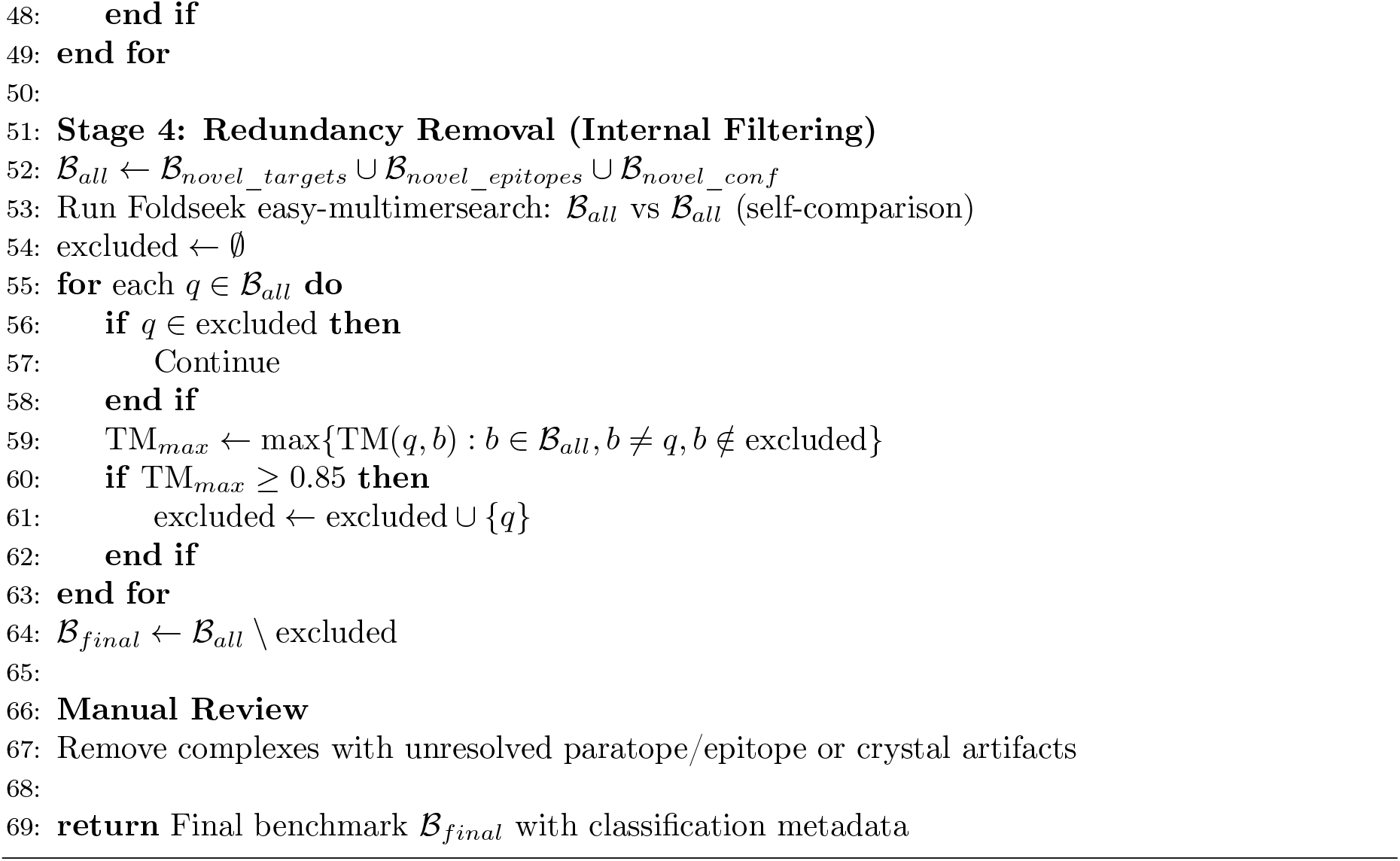

## S4 DockQ Calculation Methodology for Multi-Chain Epitopes

## S4.1 Overview

Accurate evaluation of antibody–antigen and NANOBODY^®^ VHH–antigen complex predictions requires careful handling of multi-chain epitopes, where the binding interface may span multiple antigen chains or protein subunits. Applying DockQ (Mirabello and Wallner, 2024) naively to these complexes introduces several systematic failure modes. First, DockQ’s GlobalDockQ averages over *all* pairwise chain interfaces in the input files, including antigen–antigen and heavy–light chain interfaces that are irrelevant to binding quality; this dilutes the antibody–antigen signal and biases scores downward even for correct predictions. Second, for homo-oligomeric antigens the score depends on which chain labeling is used, and the naïve label-matched evaluation may penalize a correct prediction simply because the predictor assigned chain identifiers differently from the reference. Third, when the epitope spans two antigen subunits, per-chain evaluation fragments the composite interface: a prediction that correctly docks the antibody at the subunit junction may score poorly against each chain individually while capturing the true binding mode. Finally, framework contacts from non-CDR antibody residues and crystal-packing neighbours can spuriously elevate the apparent contact count on irrelevant antigen chains, leading to incorrect multi-chain classification.

DockQ v2 provides a `–mapping` flag that can restrict evaluation to user-specified chain pairs, which partially addresses the chain-selection and label-ambiguity problems. However, it requires the correct mapping to be specified *a priori*, evaluates each chain pair as a separate pairwise interface, and provides no mechanism to merge multiple antigen chains into a composite epitope entity. It is therefore insufficient for automated, large-scale benchmarking over a heterogeneous database where epitope composition is not known in advance.

### Algorithm S3

DockQ Calculation for Multi-Chain Epitopes

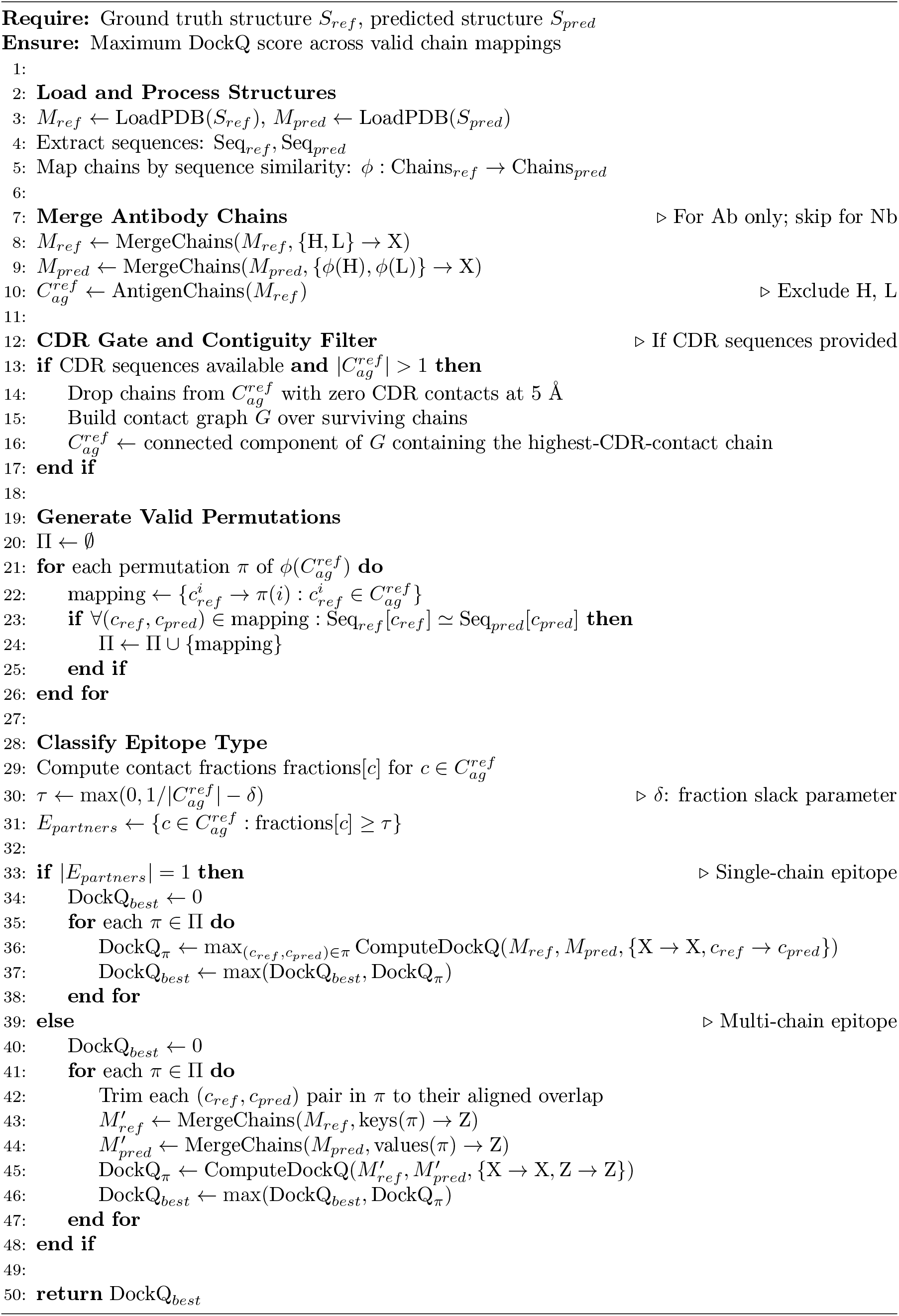

We developed a methodology that addresses each of the above issues: (1) gating antigen chains by CDR contact to exclude non-paratope and crystal-packing contributions, (2) requiring contacted chains to form a physically contiguous group, (3) classifying epitopes as single- or multi-chain based on contact fractions, and (4) exhaustively testing all valid chain permutations to handle inconsistent labeling. Algorithm S3 presents the complete workflow.

### S4.2 CDR Gate and Contiguity Filter

When CDR sequences are available, antigen chains with zero CDR contacts are dropped before the fraction-based classification. This prevents framework-only contacts from incorrectly inflating the partner count.

Among CDR-contacted chains, we additionally require physical contiguity: chains must form a connected contact graph (any atom within 5 Å). The connected component containing the chain with the most CDR contacts is retained. Disconnected chains—which may reflect crystal-packing neighbors or incidental grazing contacts—are excluded. This filter makes the epitope classification robust to the choice of the fraction slack parameter *δ* (Section S4.3). We note that the benchmark curation pipeline described in Section S3 applies equivalent criteria during complex selection, so benchmark complexes already satisfied these conditions by construction.

### S4.3 Contact-Based Epitope Classification

For each complex, we compute the fraction of total antibody interface contacts contributed by each antigen chain:

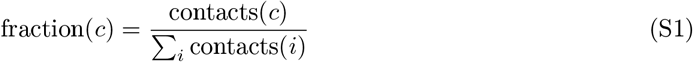

where contacts are antigen residues with at least one atom within 5 Å of the antibody or NANOB-ODY^®^ VHH. A chain is classified as an epitope partner if its fraction exceeds

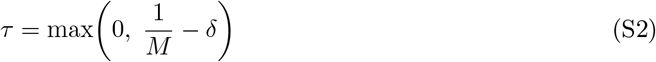

where *M* is the number of antigen chains and *δ* is a slack parameter. We evaluated *δ* ∈ {0.05, 0.15, 0.25, 0.50} and found that with the CDR gate and contiguity filter applied, classification is stable across this range; we adopt *δ* = 0.15 as a balance between conservatively requiring near-equal contact distributions and correctly capturing mildly imbalanced genuine epitopes (e.g., a minority chain contributing ∼38% of contacts).

### S4.4 Chain Permutation Handling

Chain identifiers may differ between prediction and reference files, and homo-oligomeric antigens introduce inherent labeling ambiguity. We handle this by first mapping chains via sequence similarity (≥95% identity over the aligned overlap), then enumerating all sequence-compatible permutations of the predicted antigen chains. DockQ is computed for each valid permutation and the maximum is reported. For antigens with distinct chain sequences, typically only 1–2 permutations are valid; for homo-*M* -mers, up to *M* ! are valid but this is feasible for the *M* ≤ 3 cases that constitute the majority of our benchmark.

### S4.5 Single-vs. Multi-Chain Evaluation

#### Single-chain epitopes ( |*E*_*partners*_| = 1)

DockQ is computed independently for each ref–pred antigen chain pair across all valid permutations, and the maximum is reported:

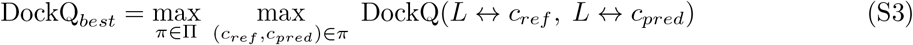

where *L* denotes the merged antibody Fv (VH-VL) or the NANOBODY^®^ VHH.

#### Multi-chain epitopes ( |*E*_*partners*_| *>* 1)

All epitope partner chains are merged into a single composite antigen entity (chain Z) before DockQ is computed. For chains of unequal length— common with near-identical homo-oligomer subunits of different lengths—chains are first trimmed to their pairwise-aligned overlap to eliminate boundary artefacts. The final score is the maximum across all valid permutations:

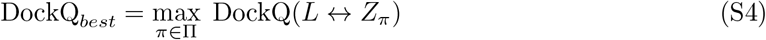

This merging strategy correctly evaluates the complete composite epitope geometry rather than fragmenting it into separate pairwise scores.

### S4.6 Antibody vs. NANOBODY^®^ VHH Handling

For antibodies, the VH and VL chains are merged into a single ligand entity prior to DockQ calculation, reflecting the fact that the paratope spans both chains. For NANOBODY^®^ VHHs (VHH only), the single heavy chain is used directly without merging.

**Figure S8:**
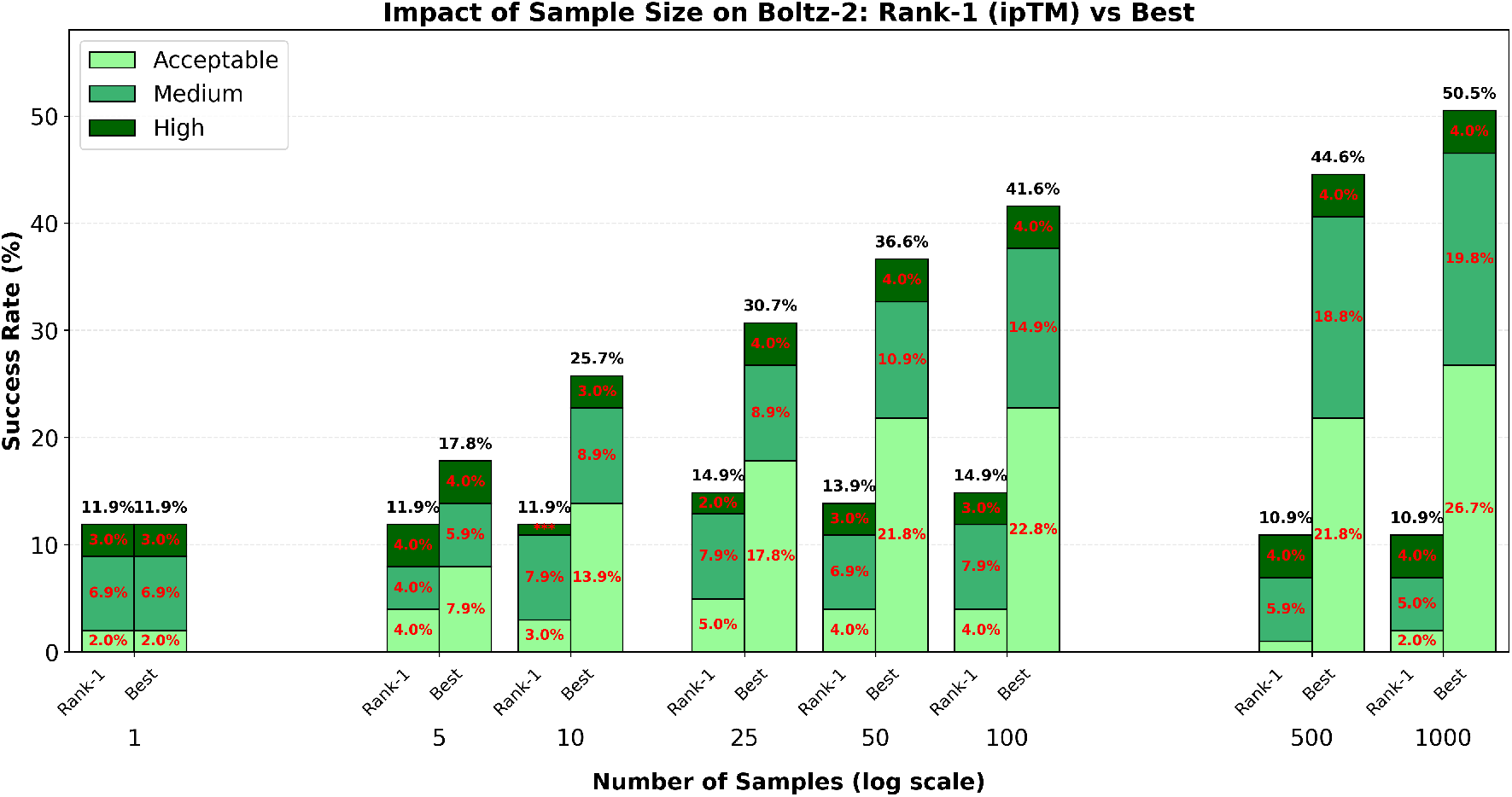
Impact of sample size on Boltz-2 performance: ipTM-based ranking vs. oracle selection. We evaluated Boltz-2’s ability to predict NANOBODY^®^ VHH-antigen complexes (N=101) using 1-1000 samples per case. For each sample size, we compared two selection strategies: **Rank-1** (selecting the prediction with highest ipTM score, as would be done in practice) versus **Best** (oracle selection of the prediction with highest DockQ score). The substantial gap between Rank-1 and Best performance reveals that while increased sampling improves the likelihood of generating accurate binding poses, current confidence scoring mechanisms fail to reliably identify them—even with 1000 samples, Rank-1 achieves only 10.9% success compared to 50.5% under oracle selection. These results demonstrate that substantial performance gains could be achieved through improved ranking mechanisms and simply generating more samples does not help.

